# eIF6 rebinding dynamically couples ribosome maturation and translation

**DOI:** 10.1101/2021.09.06.459071

**Authors:** Pekka Jaako, Alexandre Faille, Shengjiang Tan, Chi C. Wong, Norberto Escudero-Urquijo, Pablo Castro-Hartmann, Penny Wright, Christine Hilcenko, David J. Adams, Alan J. Warren

## Abstract

Protein synthesis is a cyclical process consisting of translation initiation, elongation, termination and ribosome recycling. The release factors SBDS and EFL1 (both mutated in the leukaemia predisposition disorder Shwachman-Diamond syndrome) license entry of nascent 60S ribosomal subunits into active translation by evicting the anti-association factor eIF6 from the 60S intersubunit face. Here, we show that in mammalian cells, eIF6 holds all free cytoplasmic 60S subunits in a translationally inactive state and that SBDS and EFL1 are the minimal components required to recycle these 60S subunits back into additional rounds of translation by evicting eIF6. Increasing the dose of eIF6 in mice *in vivo* impairs terminal erythropoiesis by sequestering post-termination 60S subunits in the cytoplasm, disrupting subunit joining and attenuating global protein synthesis. Our data reveal that ribosome maturation and recycling are dynamically coupled by a mechanism that is disrupted in an inherited leukaemia predisposition disorder.

## INTRODUCTION

Every minute, a growing HeLa cell synthesises around 7500 ribosomal subunits which decode messenger RNA to make protein through the four successive steps of translation: initiation, elongation, termination and recycling. Removal of the highly conserved nucleolar shuttling factor eukaryotic initiation factor 6 (eIF6) from the intersubunit face of the nascent large 60S ribosomal subunit is essential to license its entry into translation^1^, because eIF6 sterically inhibits the large 60S ribosomal subunit from joining to the small 40S subunit to form an actively translating ribosome^2,3^. eIF6 is initially recruited to pre-60S ribosomal subunits in the nucleolus^4^. Following export of the pre-60S particles to the cytoplasm, the GTPase EFL1 (elongation factor-like 1) and its cofactor SBDS (Shwachman-Bodian-Diamond syndrome) evict eIF6 during the final step in maturation of the nascent 60S subunit^5–11^.

Disruptive variants in both SBDS^12^ and EFL1^13^ cause the inherited leukaemia predisposition disorder Shwachman-Diamond syndrome (SDS)^14^. Missense variants in eIF6 can bypass the fitness defect of yeast cells lacking the SBDS orthologue Sdo1 by reducing eIF6 binding to the 60S subunit^7^. In addition, diverse somatic genetic events including point mutations, interstitial deletion and reciprocal chromosomal translocation rescue the germline ribosome defect in SBDS-deficient haematopoietic cells either by reducing eIF6 expression or by disrupting the interaction of eIF6 with the 60S subunit^15,16^. The observation that mutations in eIF6 can rescue the defects in ribosomal subunit joining and translation initiation observed in SBDS-deficient cells^15^ raises the possibility that SBDS and EFL1 may have a more general role in translation beyond their function in nascent 60S subunit maturation. Cryo-electron microscopy (cryo-EM) studies further support this hypothesis by revealing that eIF6 is bound to 60S ribosome quality control intermediates^17,18^, suggesting that there are some contexts in which eIF6 may rebind to mature 60S ribosomal subunits *in vivo*.

In eukaryotes, translation termination begins with the recognition of a stop codon in the A site of the 80S ribosome by the release factors eRF1 and GTP-bound eRF3^19^. Peptide release is temporally coupled to splitting of the 80S ribosome into a free 60S and a 40S subunit bound to deacylated tRNA and mRNA by the essential ATP-binding cassette protein Rli1 (yeast)/ABCE1 (mammals)^20,21^. The deacylated tRNA is subsequently removed, promoting dissociation of the 40S subunit from the mRNA^20,22^. ABCE1 blocks 40S rebinding to the 60S subunit by sterically hindering the formation of an intersubunit bridge between the 60S protein uL14 and the 40S rRNA helix h44^23^. However, the possibility that eIF6 rebinding might similarly sequester post-termination recycled 60S subunits in a translationally inactive state has not been addressed. Dissociated 40S and 60S subunits may immediately re-engage in further rounds of translation initiation or alternatively, in conditions of stress, enter a reservoir of translationally inactive 80S ribosomes^24–27^, that can again be recycled in an ABCE1-dependent manner^28^. Interestingly, ribosome recycling becomes critical for ribosome homeostasis during erythroid differentiation, as the natural loss of ABCE1 limits ribosome availability and results in the accumulation of post-termination, unrecycled ribosomes in the 3’UTRs of mRNAs^29^.

Here, we test the hypothesis SBDS and EFL1 act as general eIF6 release factors to regulate post-termination 60S ribosomal subunit recycling. Using cryo-EM, we show that eIF6 binds to the majority of free cytoplasmic 60S subunits in mammals, thereby holding them in a translationally inactive state. We reveal that SBDS and EFL1 are the minimal components required to evict 60S-rebound eIF6 and recycle post-termination 60S subunits back into the actively translating pool. Consistent with the requirement for efficient ribosome recycling during erythropoiesis, graded overexpression of eIF6 in mice perturbs late steps in erythroid differentiation by sequestering free 60S subunits, blocking subunit joining and attenuating global translation. Our data support a wider role for SBDS and EFL1 as translational regulators that dynamically couple 60S subunit maturation with ribosome recycling through the release of rebound eIF6.

## RESULTS

### eIF6 holds free cytoplasmic 60S subunits in a translationally inactive state *in vivo*

We set out to test the hypothesis that eIF6 maintains free cytoplasmic 60S subunits in a translationally inactive state in primary haematopoietic cells *in vivo*. Immunoblotting of cell extracts purified from primary murine c-kit+ bone marrow cells revealed that around 14 % of the eIF6 protein co-migrated with free 60S ribosomal subunits, while the majority was distributed in the free fraction (Figure 1A). Single particle cryo-electron microscopy (cryo-EM) analysis of free 60S particles purified from primary murine c-kit+ bone marrow cells revealed that eIF6 is stably bound to the intersubunit face of at least 83% of cytoplasmic mature 60S subunits (Figure 1B).

**Figure 1.**
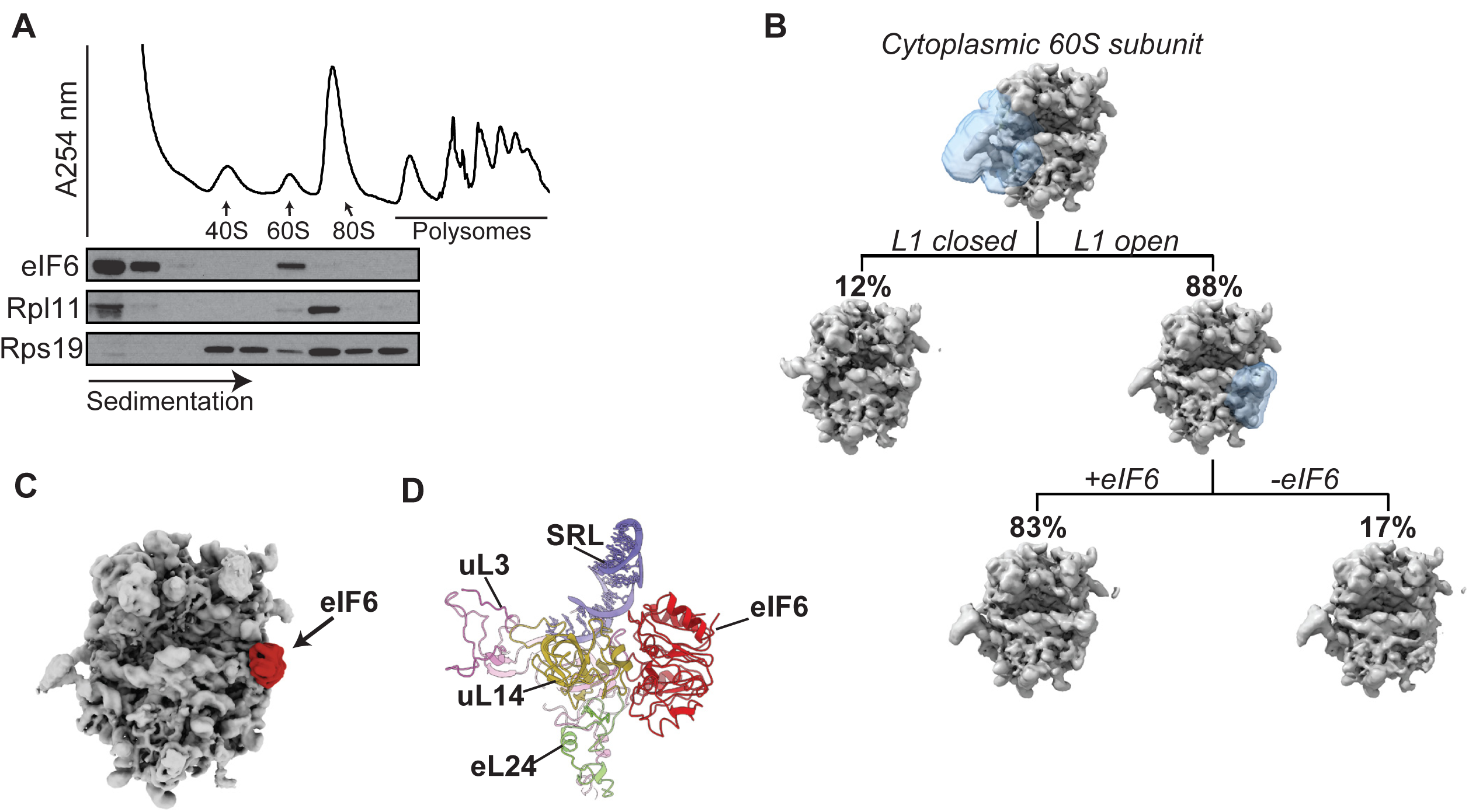
eIF6 maintains free mammalian 60S subunits in a translationally inactive state. (A) Sucrose gradient sedimentation of eIF6 in cell extracts prepared from murine c-kit+ bone marrow cells. The distribution of eIF6, Rpl11 and Rps19 was visualised by immunoblotting. (B) Cryo-EM classification scheme to quantify the frequency of eIF6-bound 60S subunits in the cytoplasm. See Material and Methods for further details. (C) eIF6 binds the intersubunit face of free cytoplasmic 60S subunits. Crown views of the cryo-EM maps of native 60S-eIF6 complexes isolated from murine c-kit+ bone marrow cells. eIF6 is highlighted by the red colour. (D) Atomic model for the murine 60S ribosomal subunit bound to human eIF6.

At an overall resolution of 3.1 Å, our cryo-EM reconstructions allowed us to build and refine atomic models of murine eIF6 bound to the 60S ribosomal subunit (Figure 1C **and** Supplementary Figure 1). Conserved from prokaryotes to human, eIF6 is a member of the pentein protein superfamily with five-fold pseudosymmetry^30^. Consistent with previous structures from yeast^3^, *Tetrahymena*^31^ and human cells^15^, murine eIF6 sterically inhibits 40S ribosomal subunit joining by binding to a conserved site on the intersubunit face of the 60S subunit involving the C terminus of uL14^32^, the sarcin-ricin loop (SRL), uL3 (residues 58–71) and the N terminus of eL24 (Figure 1D). We conclude that in primary murine haematopoietic cells, eIF6 holds all free 60S ribosomal subunits in a translationally inactive state by binding to the 60S intersubunit face. These data support the hypothesis that eIF6 must be released from the 60S ribosomal subunit to allow 80S ribosome assembly^1^. However, we were unable to discriminate nascent 60S-eIF6 complexes versus eIF6 rebound to mature 60S subunits.

### Endogenous eIF6 can rebind mature cytoplasmic 60S subunits

The ribosome quality control (RQC) pathway recognises and rescues stalled translation complexes. Following ribosome dissociation, components of the RQC complex remain bound to the 60S subunit together with eIF6^17,33^. Taken together with the finding that eIF6 is bound to virtually all mature cytoplasmic 60S ribosomal subunits, we hypothesised that during canonical translation termination (and RQC), eIF6 might rebind to mature 60S particles and require dynamic recycling by SBDS and the GTPase EFL1.

To support this hypothesis, we first tested the ability of eIF6 to rebind mature 60S particles that had been dissociated from 80S couples. Using immunoblotting, we examined the distribution of endogenous eIF6 following sucrose gradient fractionation of cell extracts prepared from c-kit+ murine bone marrow cells in 80S dissociating (2 mM Mg, 500 mM KCl) conditions. In contrast to non-dissociating conditions where eIF6 predominantly migrates in the free fraction (Figure 1A), eIF6 comigrated almost entirely with the 60S subunit (Figure 2A). Consistent with previous work^1^, we conclude that endogenous eIF6 can rebind mature cytoplasmic 60S subunits in mammalian cells.

**Figure 2.**
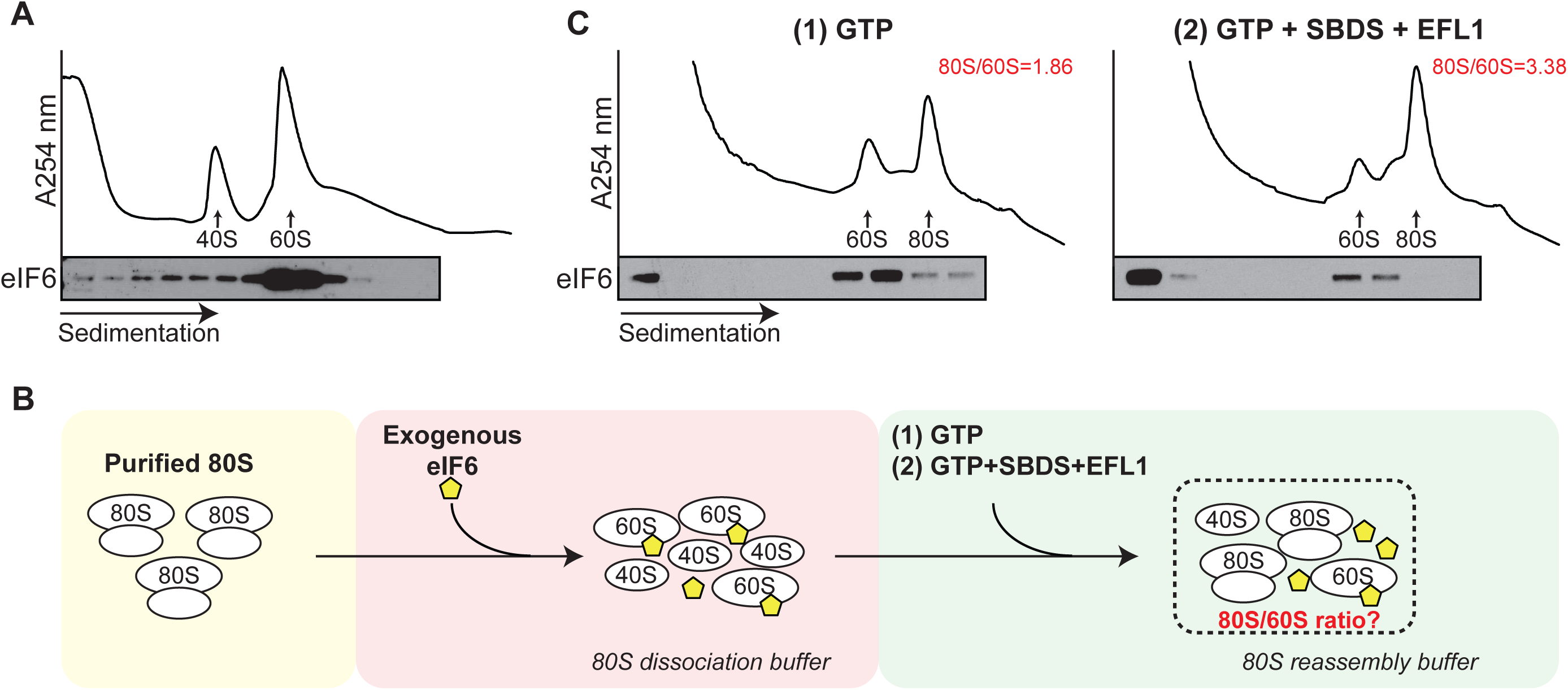
SBDS and EFL1 catalyse GTP-dependent release of rebound eIF6 from mature cytoplasmic 60S ribosomal subunits. (A) Sucrose gradient sedimentation of c-Kit+ bone marrow cell extracts (without cycloheximide) lysed in 20 mM Hepes pH 7.5, 2 mM Mg(CH_3_COO)_2_ and 500 mM KCl, and incubated for 10 min at 37 °C to allow eIF6 rebinding. eIF6 was detected by immunoblotting. (B) Schematic overview of *in vitro* eIF6 release assay. See Materials and Methods section for further details. (C) Sucrose gradient sedimentation of reconstituted eIF6 release reaction mixes. Immunoblotting was used to detect eIF6. The ratio of 80S monosomes to 60S subunits is indicated. Shown is a representative experiment out of a total of two independent experiments.

### SBDS and EFL1 are sufficient to release eIF6 rebound to 60S subunits

We next examined whether human SBDS, EFL1 and GTP are sufficient to promote the release of eIF6 rebound to mature cytoplasmic 60S subunits. We biochemically reconstituted an *ex vivo* assay that coupled eIF6 release from 60S subunits to their reassembly into 80S ribosomes by adding recombinant human SBDS and EFL1 to eIF6-loaded 60S subunits isolated from c-kit+ bone marrow cells. A schematic overview of the assay is shown in Figure 2B. As shown in the representative experiment in Figure 2C, compared with GTP alone (left panel), the addition of SBDS, EFL1 and GTP (right panel) to eIF6-loaded 60S subunits promoted redistribution of eIF6 into the free fraction of the sucrose gradient as detected by immunoblotting, with a concomitant 1.8-fold increase in 80S ribosome reassembly. We conclude that in the presence of GTP, SBDS and EFL1 are sufficient to release eIF6 that has rebound to mature 60S particles. These data provide biochemical support for the hypothesis that SBDS and EFL1 function as general release factors with dual roles in nascent 60S subunit maturation and in ribosome recycling.

### Genetic interactions between SBDS, EFL1 and eIF6

We reasoned that if eIF6 dynamically rebinds to post-termination 60S ribosomal subunits, increasing the dose of eIF6 *in vivo* would titrate out free 60S subunits to impair ribosomal subunit joining, reduce global protein synthesis and induce growth defect. Consistent with this hypothesis, ubiquitous overexpression of wild type eIF6 (but not eIF6 missense mutants identified in SDS haematopoietic cells that map to the interface with the 60S subunit) induces late larval lethality in *Drosophila*^15^. Furthermore, overexpression of SDS patient-derived eIF6 missense mutations can fully rescue the lethality of Sbds-deficient flies^15^.

To further test the *in vivo* genetic interactions between Sbds and eIF6, we depleted *Sbds* using RNAi^15^, allowing flies to develop to adult stage albeit more slowly compared with wild type controls (Figure 3A). At 29 °C, 5.4 % of *Sbds*-depleted flies develop from pupae to adults (n = 269, 3 replicates); at 25 °C, 55.5 % of pupae develop to adults (n = 276, 2 replicates). RNAi-mediated depletion of *Sbds* enhanced the growth defect induced by ubiquitous overexpression of eIF6, causing lethality at the early larval stage (Figure 3A). In the developing ommatidia, selective eIF6 overexpression induced a small, rough eye phenotype (Supplementary Figure 2) that was enhanced either by doubling the dose of eIF6 or by depleting *Sbds* or *Efl1* by RNAi (Figure 3B, Supplementary Figure 2). Selective overexpression of eIF6 in the *Drosophila* wing disc reduced global protein synthesis as measured by O-propargyl-puromycin (OP-puro) incorporation (Figure 3C). These genetic data support the hypothesis that SBDS and EFL1 function in mobilizing eIF6 that has rebound to cytoplasmic 60S ribosomal subunits *in vivo*.

**Figure 3.**
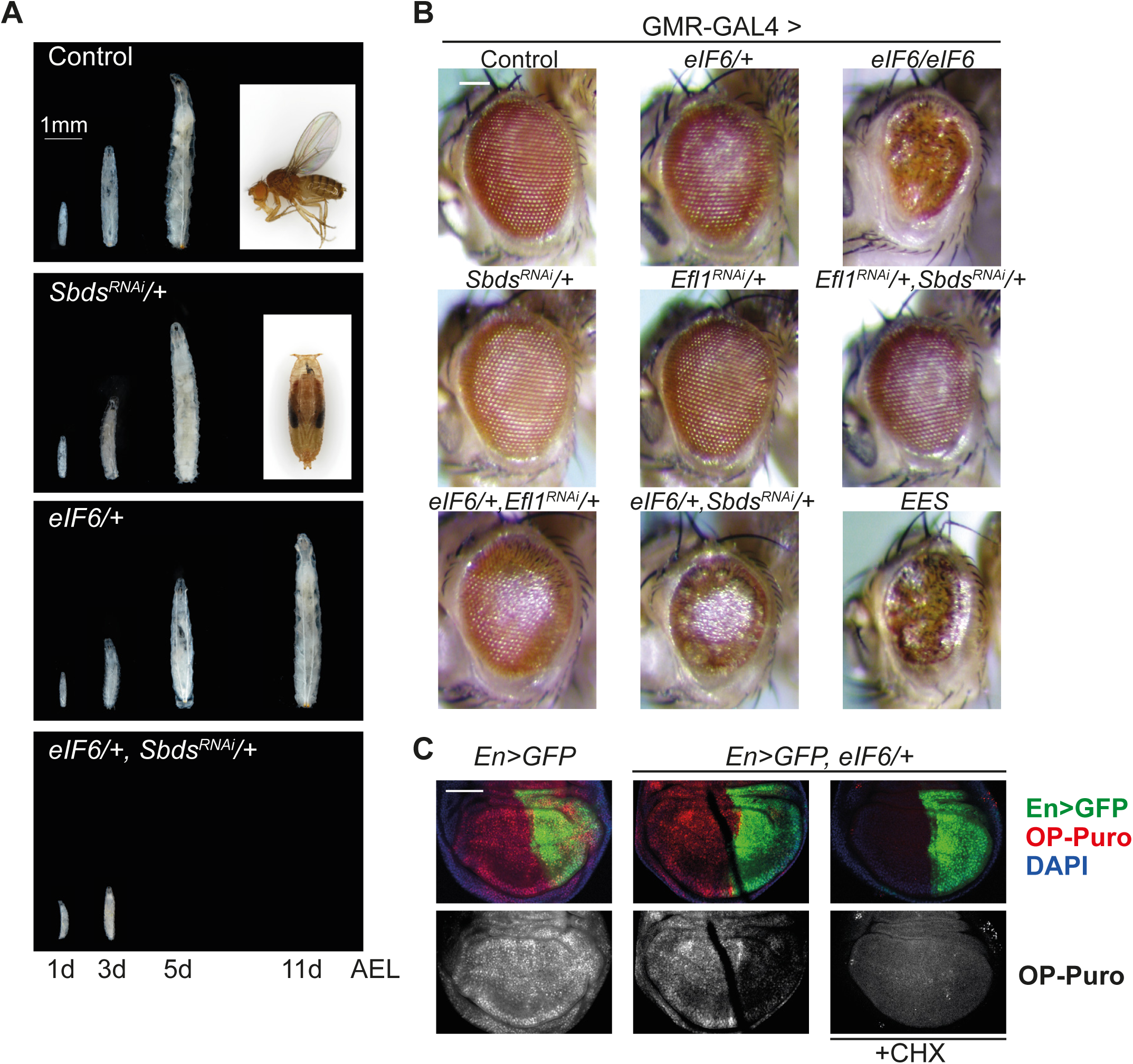
Genetic interactions between *Sbds*, *Efl1* and *eIF6*. (A) Increased eIF6 dosage enhances the growth defects of Sbds-deficient *Drosophila*. Flies were photographed at 1, 3, 5 and 11 days after egg laid (AEL). Scale bar, 1 mm. (B) Genetic interactions between *Sbds*, *Efl1* and *eIF6* in the *Drosophila* eye. Representative photomicrographs of adult eyes from flies with the indicated genotypes. *EES*, abbreviation of (*eIF6/+, Efl1^RNAi^/+, Sbds^RNAi^/+*). Scale bar, 100 µm. (C) Overexpression of eIF6 suppresses global protein synthesis in *Drosophila* wing disc cells. Third instar larval wing disc cells with the indicated genotypes were immunostained to reveal OP-Puro incorporation (red, grey). Posterior wing disc cells are marked with GFP; nucleus is blue (DAPI), scale bar: 100 µm.

### eIF6 dose-dependent inhibition of ribosomal subunit joining *in vivo*

We set out to further validate the hypothesis that eIF6 dynamically rebinds to post-termination cytoplasmic 60S ribosomal subunits by engineering a transgenic eIF6 mouse strain that permits doxycycline (Dox, tetracycline analogue)-inducible and graded overexpression of the human *EIF6* transgene by constitutively expressing the M2-reverse tetracycline transactivator (M2-rtTA) at the *Rosa26* promoter^34^ (Figures 4A, 4B). M2-rtTA is a mutant of rtTA that has increased stability, reduced background expression and improved inducibility in the presence of Dox^35^. This transgenic mouse strain exhibits widespread constitutive expression of M2-rtTA, allowing for Dox-inducible transactivation of the human *EIF6* cDNA. We adjusted the level of eIF6 overexpression by breeding animals that were heterozygous or homozygous for the *M2-rtTA* at the *Rosa26* locus and carried one or two copies of the human *EIF6* transgene at the *Col1a1* locus (Figure 4B). To evaluate the level of *EIF6* transgene expression, we treated cultured c-Kit+ bone marrow cells with Dox and performed quantitative real-time PCR to measure *EIF6* mRNA. We designed two sets of primers to distinguish endogenous mouse *Eif6* mRNA from total (endogenous + transgene) *EIF6* mRNA to verify the transgene copy number–dependent increase in total *EIF6* expression (Figure 4C). The increase in *EIF6* mRNA led to a significant increase in the level of eIF6 protein expression compared with control animals (Figure 4D).

**Figure 4.**
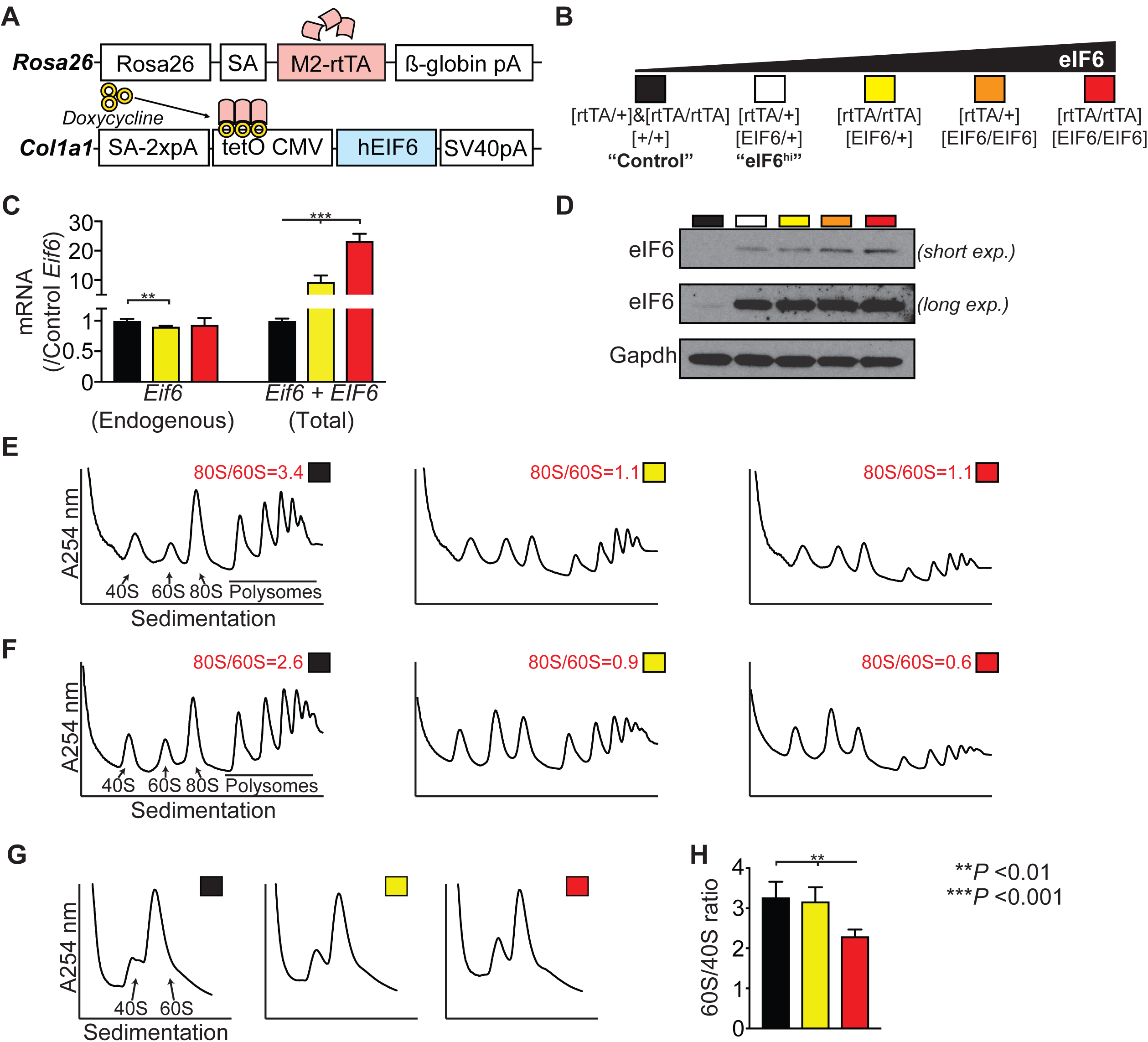
eIF6 binds post-termination 60S subunits to prevent ribosomal subunit joining. (A) Schematic overview of the transgenic Dox-inducible eIF6 overexpression system. (B) Breeding strategy for graded overexpression of eIF6, with colour coding of indicated genotypes. (C) Quantitative real-time PCR of *EIF6* transcript levels (n=3-4 per genotype). (D) eIF6 protein immunoblotting analysis in extracts from cultured c-Kit+ bone marrow cells derived from the indicated mouse strains after 24 hr of Dox induction. (E, F). Sucrose gradient sedimentation of extracts (including cycloheximide) from cultured c-Kit+ bone marrow cells derived from the indicated mouse strains. Dox induction, 24 hr. Buffers in (E) and (F) contain 50 mM or 200 mM KCl, respectively. Shown is representative of two independent experiments. (G) Sucrose gradient sedimentation of extracts prepared in absence of magnesium to dissociate 80S ribosomes and polysomes. (H) Quantification of the 60S:40S subunit ratios shown in (G) (n=3 per genotype). Student’s *t* test was used to determine statistical significance, and two-tailed *P* values are shown. Error bars represent standard deviation.

Next, we assessed the impact of increasing doses of eIF6 on ribosome assembly *in vivo* by fractionating cell extracts in the presence of cycloheximide from Dox-treated c-Kit+ bone marrow cells by sucrose gradient sedimentation. Depending on the level of overexpression, the increased dose of eIF6 promoted a reduction in the 80S:60S ratio, consistent with a subunit-joining defect (Figure 4E). Parallel experiments using high salt buffer to specifically dissociate inactive mRNA-free 80S monosomes^36^, further highlighted the eIF6 dose-dependent reduction in actively translating 80S ribosomes (Figure 4F). Finally, by using a magnesium-free buffer system, we observed that the ratio of 60S to 40S subunits was preserved with an intermediate dose of eIF6 (Figure 4G). Although higher eIF6 overexpression resulted in a relative decrease in 60S subunits (Figure 4H), this is likely to be a secondary consequence of the profound reduction in global protein synthesis. We conclude that graded eIF6 overexpression induces a dose-dependent ribosomal subunit joining defect *in vivo*. Importantly, the observed ribosomal subunit joining defect upon eIF6 overexpression closely mimics the subunit joining defect caused by eIF6 retention on the 60S subunit that is observed in *Sbds*- or *Efl1* deficient mice or patient-derived lymphoblasts^8,9,13,37^. Taken together with our genetic data in *Drosophila*, we propose that the most logical interpretation of these findings is that eIF6 rebinds to post-termination recycling 60S subunits from which it is dynamically recycled by SBDS and EFL1. These data support the hypothesis that SBDS and EFL1 translationally activate nascent 60S subunits and in addition act as general eIF6 release factors that dynamically recycle eIF6-bound post-termination 60S subunits back into additional rounds of translation.

### Terminal erythroid differentiation is sensitive to eIF6 dosage

We reasoned that during mammalian haematopoiesis, the erythroid lineage might be particularly sensitive to an increased dose of eIF6 and aberrant ribosome homeostasis due to the increased dependence of terminal erythroid differentiation on ribosome recycling as a consequence of natural loss of the ribosome recycling factor ABCE1^29^. To test this hypothesis, we induced eIF6 overexpression *in vivo* in transgenic mice.

Detailed analysis of mice carrying two copies of the *M2-rtTA* transgene and either one or two copies of the *EIF6* transgene was precluded because of the rapid weight loss induced in these animals. By contrast, mice that were heterozygous for both transgenes (*M2-rtTA/+*; *EIF6/+*, herein called eIF6^hi^ mice) did not lose weight acutely in response to Dox administration (Supplementary Figure 3). We therefore restricted our analysis to eIF6^hi^ mice.

Immature (lineage-, Sca-1+, c-Kit+; LSK), myeloid (preGM/GMP) and erythroid (preCFU-E/CFU-E) haematopoietic progenitor cells isolated from Dox-treated eIF6^hi^ mice showed a 2-4 fold increase in *EIF6* mRNA (Supplementary Figure 4A), while sucrose gradient sedimentation analysis of extracts from cultured c-Kit+ bone marrow cells showed accumulation of free 40S and 60S subunits compared with control (Supplementary Figure 4B). Immunoblotting revealed a robust increase in eIF6 protein across the gradient (Supplementary Figure 4C). Compared with controls, the overexpressed eIF6 protein predominantly accumulated in the cytoplasm of freshly isolated bone marrow cells in eIF6^hi^ mice (Supplementary Figure 4D).

After two weeks of Dox administration, eIF6^hi^ mice developed persistent macrocytic anaemia with a significant reduction in the reticulocyte count compared with controls (Figure 5A **and** 5B). While the platelet count increased, the total white blood cell count was unaffected (Supplementary Figure 5). Histological examination of the bone marrow revealed erythroid hyperplasia in eIF6^hi^ mice, with an increased frequency of erythroid precursors compared with controls (Supplementary Figure 6A). In addition, the spleen was enlarged in eIF6^hi^ mice (Supplementary Figure 6B), due to marked expansion of erythroid precursors (Supplementary Figure 6C).

**Figure 5.**
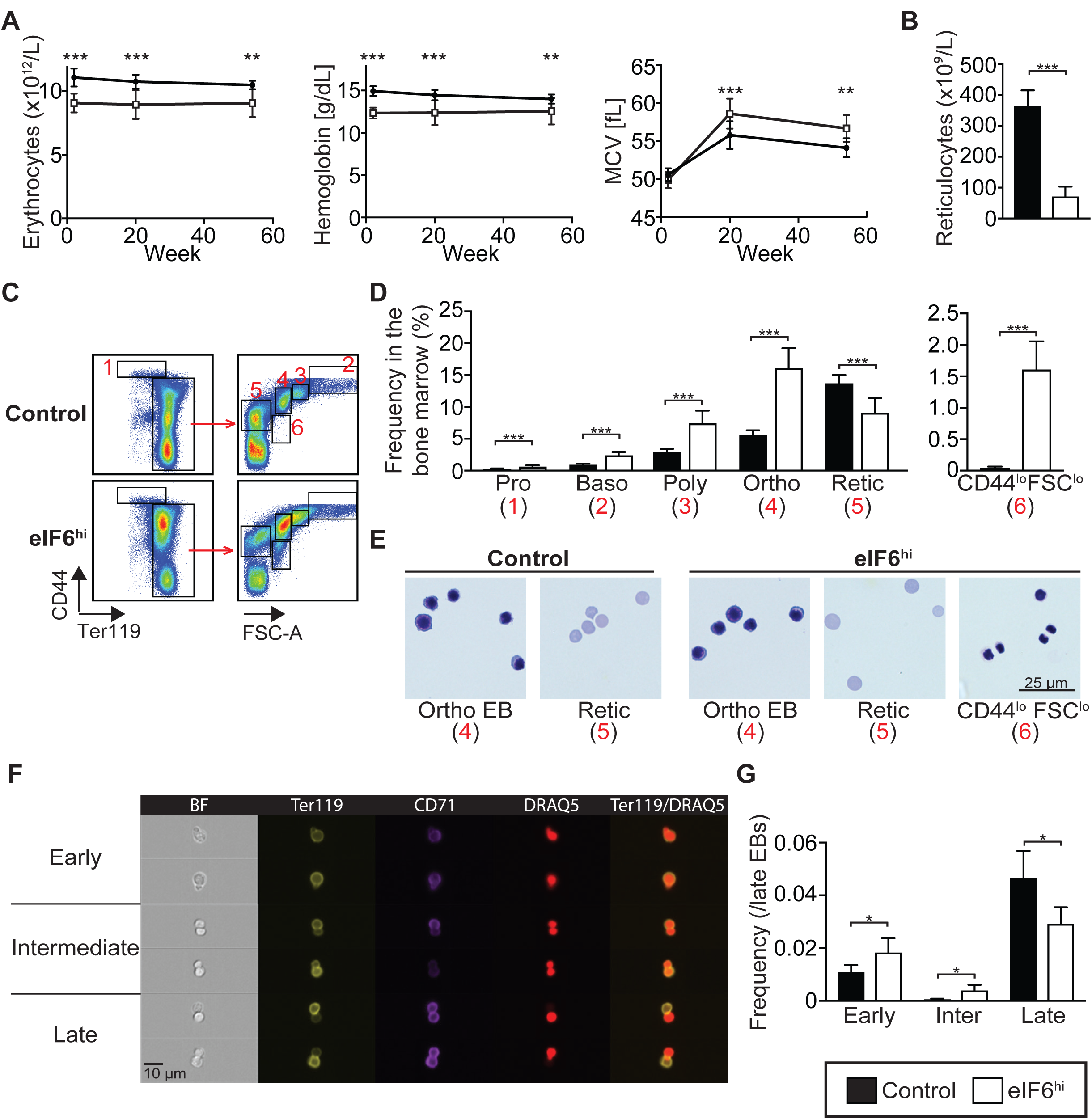
Increased eIF6 dosage impairs erythroblast enucleation in mice. (A) Increased eIF6 dosage causes macrocytic anemia. Haematological parameters including hemoglobin concentration, erythrocyte count and mean corpuscular volume (MCV) (n=12-15 per genotype) are shown over the indicated time-course of Dox induction for eIF6^hi^ mice versus control. (B) Reticulocyte counts (n=3 per genotype). (C) Representative flow cytometry analysis of erythroid precursors in control versus eIF6^hi^ bone marrow. Gated populations are designated 1-6 in red. (D) Frequency of erythroid precursors in the bone marrow (n=7 per genotype), corresponding to gated populations 1-6 in the flow cytometry analysis. Pro, proerythroblast; Baso, basophilic erythroblast; Poly, polychromatic erythroblast; Ortho, orthochromatic erythroblast; Retic, reticulocyte. (E) Morphology of erythroid precursors, corresponding to populations 4-6 by flow cytometry. (F) Representative images of enucleating erythroblasts, defined by Amnis ImageStream IDEAS gating strategy, shown in Supplementary Figure 9. (G) Frequencies of enucleating erythroblasts within the late erythroblast population (corresponding to gate 6 in IDEAS gating strategy), in the bone marrow (n=4 per genotype) after 2 weeks of Dox administration. Student’s *t* test was used to determine statistical significance, and two-tailed *P* values are shown. Error bars represent standard deviation.

To further characterise haematopoiesis in the eIF6^hi^ mice, we analysed bone marrow cells by flow cytometry^38,39^, using the gating strategy shown schematically in Supplementary Figure 7. The eIF6^hi^ mice showed no significant differences in overall bone marrow cellularity relative to controls (Supplementary Figure 8A). Although the frequency of myeloid and multipotent progenitors (preGM and MPPs) decreased, the frequency of erythroid progenitors (preCFU-E and CFU-E) (Supplementary Figure 8B) and precursor cells (Figure 5C-E) was significantly increased. A similar increase in the frequency of erythroid precursors was detected by flow cytometry in the spleen (Supplementary Figure 8C). Within the bone marrow, we identified an abnormal population of orthochromatic erythroblast-like cells (CD44^lo^ FSC^lo^) containing a highly condensed nucleus and low cytoplasmic volume (Figure 5C-E).

We hypothesised that an increased dose of eIF6 might impair erythroblast enucleation during the terminal steps of erythroid differentiation, promoting the accumulation of orthochromatic erythroblast-like cells, but reducing the numbers of reticulocytes. To test this, we applied Amnis ImageStream technology^40,41^ to visualise active nuclear extrusion by bone marrow erythroblasts, dividing the process into early, intermediate and late stages (Figure 5F **and** Supplementary Figure 9). Compared with controls, in Dox-treated eIF6^hi^ mice we classified more erythroblasts in the early or intermediate stages of enucleation compared with late steps (Figure 5G). We conclude that an increased dose of eIF6 impairs terminal enucleation of orthochromatic erythroblasts *in vivo*.

We next set out to determine whether the eIF6-dependent erythroid differentiation defect was intrinsic to eIF6^hi^ haematopoietic cells. Consistent with this hypothesis, *ex vivo* differentiation of CFU-Es/proerythroblasts isolated from Dox-treated eIF6^hi^ mice recapitulated the eIF6-dependent defect in terminal erythropoiesis (Supplementary Figure 8D). Furthermore, non-competitive transplantation of bone marrow cells from eIF6^hi^ mice into lethally irradiated wild type congenic recipients also recapitulated the eIF6 dose-dependent haematopoietic abnormalities (Supplementary Figure 10). Taken together, our data indicate that the terminal erythroid maturation defects are intrinsic to eIF6^hi^ haematopoietic cells.

### Attenuated protein synthesis impairs terminal erythroblast enucleation

We hypothesised that increasing the dose of eIF6 would alter ribosome homeostasis during erythropoiesis by shifting the equilibrium towards ribosomal subunit dissociation, thereby attenuating protein synthesis. To test this, we quantified the rate of global protein synthesis in erythroid cells *in vivo* by measuring OP-puro incorporation. Indeed, compared with controls, we observed a significant decrease in OP-puro incorporation in late poly- and orthochromatic erythroid precursors from Dox-treated eIF6^hi^ mice (Figure 6A). These data demonstrate that eIF6 overexpression impairs terminal erythroid differentiation by a mechanism that directly or indirectly attenuates protein synthesis.

**Figure 6.**
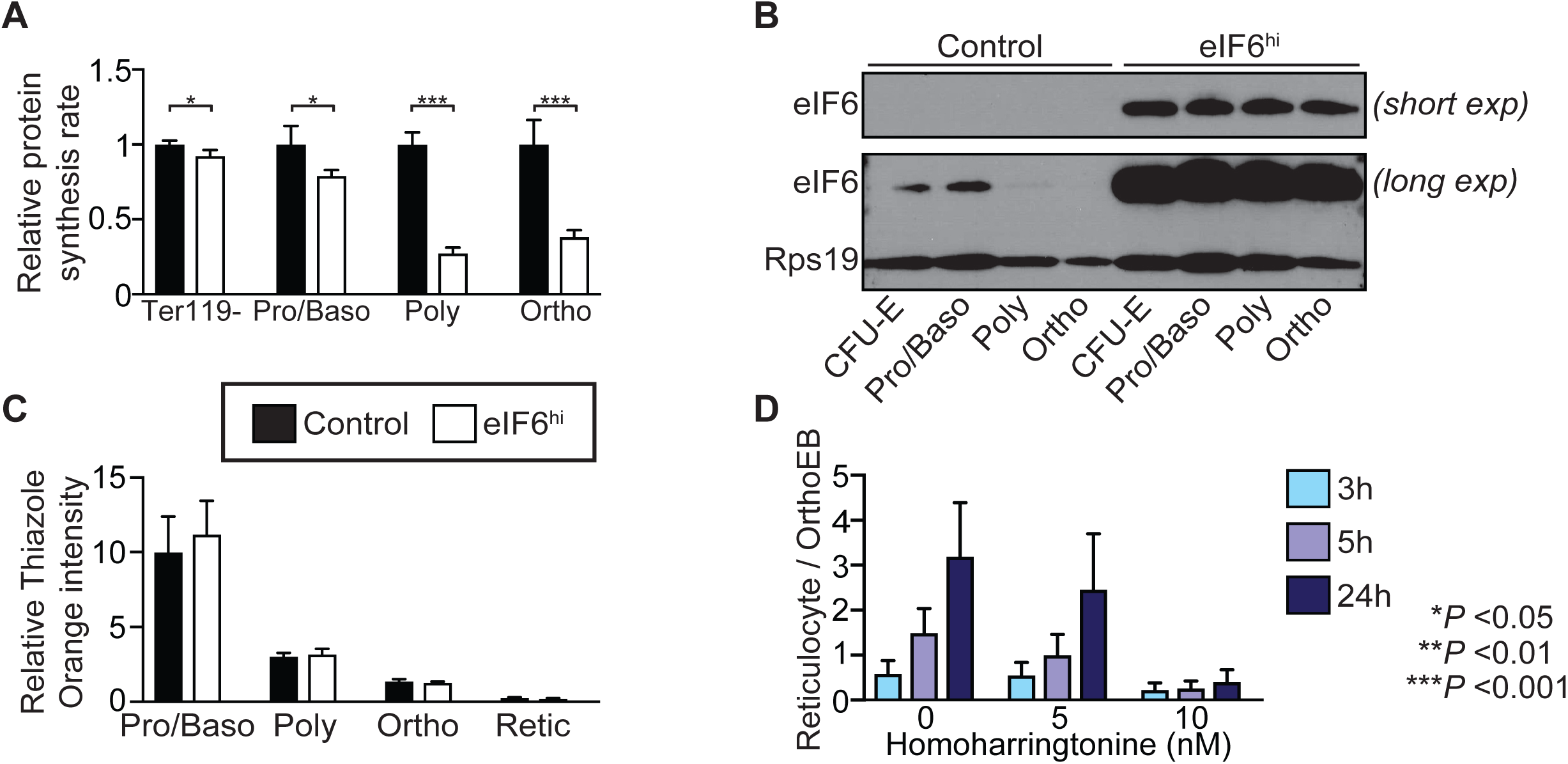
Increased eIF6 dosage impairs erythroblast enucleation by attenuating protein synthesis. (A) OP-Puro incorporation in the indicated bone marrow cells *in vivo* after two weeks of Dox administration (n=3-4 per genotype). Median fluorescence intensities were normalised against the respective control cell populations. (B) Expression of eIF6 in CFU-E erythroid progenitor cells and erythroid precursors *in vivo* after two weeks of Dox treatment. Immunoblots are shown for eIF6 and Rps19 using extracts generated from identical numbers of the indicated bone marrow cells. Shown is representative of two independent experiments. CFU-E progenitor cells are defined as CD71+ TER-119-bone marrow cells. (C) Total cellular nucleic acid content *in vivo* during terminal erythroid differentiation. Freshly isolated bone marrow cells (n=4 per genotype) were stained with thiazole orange. Thiazole orange intensities are shown relative to CD44+ TER-119- non-erythroid bone marrow cells. (D) Enucleation of FACS-purified wild type orthochromatic erythroblasts in culture after 3, 5 or 24 hr treatment with homoharringtonine (n=3). Enucleation efficiency is expressed as the ratio of reticulocytes to orthochromatic erythroblasts. Student’s *t* test was used to determine statistical significance, and two-tailed *P* values are shown. Error bars represent standard deviation.

We reasoned that the reduced rate of protein synthesis in late erythroblasts from Dox-treated eIF6^hi^ mice likely reflects altered ribosome homeostasis as a consequence of an increase in the relative ratio of eIF6 to ribosomes during terminal erythroid differentiation. To test this hypothesis, we sorted identical numbers of erythroid progenitor and precursor cells from Dox-treated mice and performed immunoblotting to visualise eIF6 and Rps19 (as a marker for cellular ribosome levels). In control mice, the levels of eIF6 and Rps19 peaked in early erythroblasts and progressively declined during terminal erythroid differentiation (Figure 6B). By contrast, erythroblasts in Dox-induced eIF6^hi^ mice exhibited sustained high levels of eIF6 (Figure 6B). The relative intensity of thiazole orange staining (correlating with cellular ribosomal RNA content) of freshly isolated erythroblasts was consistent with a progressive decline in cellular ribosome levels during terminal erythroid maturation (Figure 6C). Taken together, these results indicate that an increased dose of eIF6 relative to ribosomal subunits is sustained in the eIF6^hi^ mice throughout erythropoiesis. Erythroid differentiation is likely susceptible to increased eIF6 dosage due to the combined shutdown in new ribosome synthesis in early erythroblasts^42^ together with the loss of effective ribosome recycling through natural loss of the ribosome recycling factor ABCE1 during terminal differentiation^29^. We propose that the increased dose of eIF6 titrates out recycled post-termination 60S subunits during late erythroid differentiation to push the equilibrium in favour of ribosomal subunit dissociation, impaired translation initiation and attenuated protein synthesis. Finally, consistent with the impact of eIF6 overexpression on terminal erythroid differentiation, inhibition of protein synthesis with the translational elongation inhibitor homoharringtonine in prospectively isolated wild-type orthochromatic erythroblasts recapitulated the erythroblast enucleation defect observed in eIF6^hi^ mice (Figure 6D).

## DISCUSSION

In this study, we have identified a critical role in the regulation of translation initiation for the SBDS and EFL1 proteins through their role as general eIF6 release factors. Using cryo-EM, we provide direct evidence that eIF6 holds virtually all free cytoplasmic 60S subunits in mammalian cells in a translationally inactive state and show that SBDS and EFL1 are the minimal components required to recycle eIF6 that has rebound to post-termination 60S subunits. Depletion of Sbds or Efl1 exacerbates the growth defects caused by eIF6 overexpression in *Drosophila in vivo*, while eIF6 overexpression in mice causes a dose-dependent defect in ribosomal subunit joining by rebinding and titrating out post-termination 60S subunits from active translation. The observation that inactive 80S monosomes accumulate in eIF6 haploinsufficient mice^43^ also supports the hypothesis that eIF6 prevents the formation of inactive 80S monosomes by binding to post-termination 60S subunits. Taken together, our data support a role for SBDS and EFL1 in regulating ribosome homeostasis by coupling the final step in cytoplasmic 60S subunit maturation with post-termination 60S ribosomal subunit recycling through their role as general eIF6 release factors (Figure 7).

**Figure 7.**
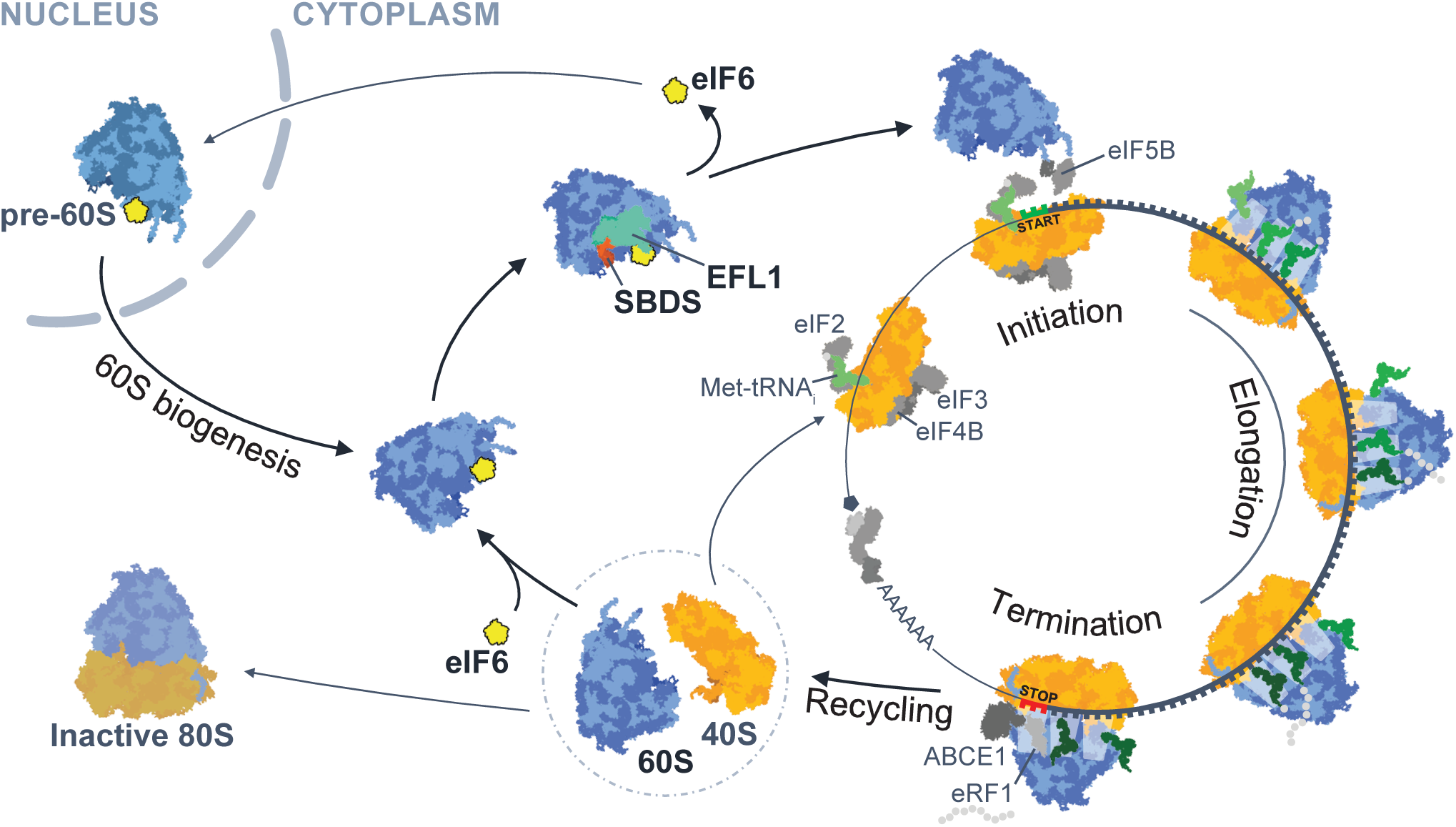
Model illustrating how dynamic rebinding of eIF6 couples ribosome maturation and translation. eIF6 functions as a ribosome anti-association factor to hold nascent pre-60S and mature post-termination 60S subunits in a translationally inactive state. SBDS and EFL1 couple nascent 60S subunit maturation and ribosome recycling by acting as general eIF6 release factors.

Translation of mRNA occurs in four steps: initiation, elongation, termination and ribosome recycling. During the normal translation cycle, once the ribosome reaches the stop codon of the mRNA, eRF1 and eRF3 recognise the stop codon and trigger hydrolysis of the nascent chain. Upon dissociation of eRF3, 80S ribosomes are recycled by recruitment of the ATPase ABCE1 to regenerate free 40S and 60S subunits^20^. This process maintains ribosome homeostasis by promoting additional rounds of translation initiation. Following 80S ribosome dissociation, the free 60S subunit may re-enter a new round of translation by binding a 48S pre-initiation complex to form an elongation competent 80S. Alternatively, it may bind an empty 40S subunit to form a vacant mRNA-free 80S monosome. A third possibility is that post-termination 60S subunits bind eIF6 to maintain the cytoplasmic pool of free ribosomal subunits in a translationally inactive state.

This begs the question of how translationally inactive eIF6-bound 60S subunits are recycled back into active translation. Genetically, depletion of SBDS and EFL1 reduces global protein synthesis due to the defect in ribosomal subunit joining caused by eIF6 retention on the intersubunit face of the 60S subunit^7,13,15^. While SBDS and EFL1 are known to release eIF6 during the final cytoplasmic step in nascent 60S maturation, the marked reduction in protein synthesis in SBDS and EFL1-deficient cells suggested to us that these factors may have a broader role as general release factors that liberate rebound eIF6 in a number of different contexts such as during post-termination ribosome recycling.

Although eIF6 was shown to bind to free 60S subunits by immunoblotting of mammalian cell extracts fractionated by sucrose gradient sedimentation^1^, the stoichiometry of this interaction *in vivo* remained unclear. In this study, we show that increasing the dose of eIF6 *in vivo* alters ribosome homeostasis by sequestering all free post-termination cytoplasmic 60S subunits, impairing ribosomal subunit joining and reducing 80S assembly. The subunit joining defect induced by eIF6 overexpression mimics the consequences of SBDS or EFL1 deficiency in SDS patient cells, *Dictyostelium*, mice and zebrafish^8,9,13,37^ and is exacerbated by concomitant depletion of either SBDS or EFL1. Our data therefore suggest that increasing the dose of eIF6 alters ribosome homeostasis by exceeding the capacity of endogenous SBDS and EFL1 to evict eIF6 from dynamically recycling 60S subunits.

Our findings suggest that the inability to dynamically upregulate recycling of post-termination ribosomes back into active translation at key time points during development may be a critical facet of SDS pathogenesis. This is exemplified by the defect in erythroid differentiation we observed in mice expressing an increased dosage of eIF6. Our model provides a more satisfactory explanation of why diverse mosaic somatic genetic events, including point mutations, interstitial deletion and reciprocal chromosomal translocation involving *EIF6* may confer a selective advantage in SBDS-deficient haematopoietic cells^15^ by disrupting the expression of eIF6 or its interaction with cytoplasmic (but not nuclear) 60S subunits, thus still preserving ribosome biogenesis. Indeed, SDS-related somatic *EIF6* missense mutations that reduce eIF6 dosage or binding to cytoplasmic 60S subunits suppress the ribosome assembly and protein synthesis defects across multiple SBDS-deficient species including yeast, *Dictyostelium*, *Drosophila* and human cells^15^. Taken together, these genetic and biochemical data support a major role for SBDS and EFL1 in regulating cytoplasmic ribosome homeostasis and translational control.

As our transgenic mice overexpressing eIF6 recapitulate the defect in ribosome assembly observed in SDS, this model may provide a tool to further dissect SDS pathogenesis. Similar to germline depletion of Sbds or Efl1 in mice^13,37,44,45^, high doses of eIF6 are not systemically tolerated. However, future studies combining the *EIF6* transgene with tissue-specific tetracycline transactivator mouse strains will bypass this limitation, harnessing the full potential of this model. Finally, our inducible eIF6 transgenic mouse model may find utility in the development of therapeutic strategies to restore cytoplasmic ribosome homeostasis in SDS by modulating the rebinding of eIF6 to cytoplasmic 60S subunits.

## MATERIALS AND METHODS

### Generation of transgenic eIF6 mouse strain

Gibson assembly was used to clone a full-length human *EIF6* cDNA containing Kozak sequence (5’-ATCACG-3’) into the EcoRI site of pBS31 vector, which was in turn used to target the KH2 embryonic stem (ES) cell line^34^. The engineered ES cells were injected into E3.5 C57BL/6 blastocysts to generate chimeric mice. Mice were backcrossed into the C57BL/6 background for at least three generations. PCR was used to genotype the *Rosa26* locus (5’-AAAGTCGCTCTGAGTTGTTAT-3’; 5’-GCGAAGAGTTTGTCCTCAACC-3’; 5’-GGAGCGGGAGAAATGGATATG-3’; WT product: 600 bp; Insert product: 300 bp) and the *Col1a1* locus (5’-TCCCTCACTTCTCATCCAGATATT-3’; 5’-AGTCTTGGATACTCCGTGACCATA-3’; 5’-GGACAGGATAAGTATGACATCATCAA-3’; WT product: 1092 bp; Insert product: 455 bp). The *EIF6* transgene was induced *in vivo* by administering Dox in the food (ssniff-Spezialdiäten GmbH; 2000 mg/kg). Mice were maintained in specific pathogen-free conditions and all procedures were performed according to the United Kingdom Home Office regulations. All experiments were performed using adult (8-12 weeks old) female and male mice with littermate controls.

### Peripheral blood analysis

Peripheral blood was collected from the tail vein into Microvette® 500 K3E tubes (Sarstedt) and cellularity analysed using a Woodley ABC blood counter.

### Histopathology

Organs for histopathological analysis were fixed in 4 % formaldehyde (Genta Medical, UK) followed by paraffin embedding and sectioning. Sections were stained with Hematoxylin-Eosin (Merck) for microscopic examination. FACS-purified erythroid precursors were transferred onto slides using a cytospin centrifuge and stained with May-Grünwald and Giemsa solutions (Merck). Morphological examination was performed using AxioImager Z2 Upright Wide-field Microscope (Zeiss).

### Flow cytometry

We isolated bone marrow cells by crushing hips, femurs and tibias in PBS (Thermo Fisher Scientific) supplemented with foetal calf serum (FCS; 2 %; Thermo Fisher Scientific) and EDTA (2 mM; Thermo Fisher Scientific). Isolated cells were filtered through a 70 µm cell strainer (Thermo Fisher Scientific). Antibody labelling was performed in PBS (+2 % FCS) for 30 min on ice. Antibodies are listed in the Supplementary Table 1. Erythrocytes were removed from peripheral blood by Dextran sedimentation (2 % in PBS; Merck) and ACK lysis buffer (Thermo Fisher Scientific) before antibody labelling. Experiments were performed using FACSARIA III cell sorter (BD Biosciences) and LSRFortessa flow cytometer (BD Biosciences), and analysed using FlowJo software (Tree Star, v10.1r7).

### Imaging flow cytometry

Sample preparation was performed as previously described^41^. Briefly, 10 × 10^6^ unfractionated bone marrow cells were fixed using formaldehyde (4 %; Alfa Aesar) for 15 min at room temperature. Following two washes with PBS, the cell pellet was cooled on ice for 15 min, and permeabilised using ice-cold acetone (a cycle of 50%-100%-50%). Following a wash with PBS (+2 % FCS), cells were stained for surface markers. Finally, 10 × 10^6^ cells were resuspended in 100 µL PBS supplemented with DRAQ5 (2.5 µM; BioLegend), with acquisition performed on an ImageStream®X Mark II Imaging Flow Cytometer (Merck) using a 40 × objective lens. Approximately 50 000 events per sample were collected, and data analysis was performed using the associated Image Data Exploration and Analysis software (IDEAS; v.6.2; Merck).

### Cell isolation and culture

c-Kit+ bone marrow cells were enriched using CD117 MicroBeads and MACS separation columns (Miltenyi Biotec), and cultured in OptiMEM I reduced Serum Media (Thermo Fisher Scientific), supplemented with FCS (10 %), penicillin/streptomycin (P/S, Life Technologies), β-mercaptoethanol (50 µM; Thermo Fisher Scientific), murine stem cell factor (mSCF; 100 ng/mL, PeproTech), murine interleukin 3 (mIL-3; 10 ng/mL, PeproTech) and murine granulocyte-colony stimulating factor (mG-CSF; 10 ng/mL, PeproTech) ± Dox (1 µg/mL; Merck). Biotinylated antibodies and Anti-Biotin MicroBeads (Miltenyi) were used for lineage depletion. *In vitro* erythroid culture was performed as previously described^46^. Briefly, 1-2.5 × 10^5^ CFU-E/proerythroblasts isolated from Dox-treated mice were seeded on fibronectin-coated (2 µg/mL; Merck) 48-well plates in Iscove’s modified Dulbecco’s medium (IMDM; Thermo Fisher Scientific) containing FCS (15 %), bovine serum albumin (BSA; 1 %; Stem Cell Technologies), mSCF (10 ng/mL), recombinant human erythropoietin (10 U/mL; Cell Signaling Technology), human recombinant insulin (100 µg/mL; Merck), recombinant human insulin-like growth factor 1 (hIGF1; 100 ng/mL; Thermo Fisher Scientific), holo-transferrin (200 µg/mL; Merck), L-glutamine (2 mM; Merck), β-mercaptoethanol (50 µM) and P/S. The following day, culture media was replaced with differentiation media consisting of IMDM, FCS (20 %), β-mercaptoethanol (50 µM), P/S and L-glutamine (2 mM). Homoharringtonine-supplemented differentiation media was used to assess the enucleation of prospectively purified orthochromatic erythroblasts.

### Transplantation assays

Non-competitive transplantations were performed by injecting 5 × 10^6^ freshly isolated unfractionated bone marrow cells in 250 µL PBS (+2 % FCS) into the tail vein of lethally irradiated (2x 500 cGY) congenic (CD45.1) wild-type recipients. Reconstituted mice were allowed to recover for two weeks before Dox administration.

### Protein synthesis rate measurement

O-propargyl-puromycin (OP-puro) labelling experiments were performed as previously described^47^. Briefly, OP-puro (50 mg/kg in 200 µL PBS; Jena Bioscience) was injected intraperitoneally and bone marrow cells were isolated after 1 hr. 3 × 10^6^ cells were fixed with formaldehyde (4 %) for 15 min at room temperature. Following two washes with PBS (+2 % FCS), cells were stained with antibodies against cell surface markers. Stained cells were permeabilised using PBS supplemented with saponin (0.1 %; Merck) and FCS (2 %). The Click reaction was performed using the Click-iT™ Plus OPP Alexa Fluor™ 488 Protein Synthesis Assay Kit according to manufacturer’s instructions (Thermo Fisher Scientific).

### Polysome profiling experiments

Equal numbers of c-Kit+ bone marrow cells were expanded in the presence of doxycycline for 24 hours and treated with cycloheximide (CHX; 100 µg/mL; Merck) for 8 min at 37 °C before harvesting by centrifugation. Cells were washed twice with ice-cold PBS supplemented with CHX (100 µg/mL) and lysed for 30 min on ice in ‘standard’ lysis buffer (20 mM Hepes pH 7.5, 50 mM KCl, 10 mM Mg(CH_3_COO)_2_, 100 µg/mL CHX, cOmplete™ EDTA-free Protease Inhibitor Cocktail (Merck), RNaseOUT recombinant ribonuclease inhibitor (200 U/mL; Thermo Fisher Scientific) and IGEPAL® CA-630 (0.5 %; Merck), Dithiothreitol (DTT; 2 mM; Merck). The lysate was cleared by centrifugation (18 000 g for 8 min at 4 °C), and loaded onto a 5-45 % (w/v) sucrose gradient (prepared in 20 mM Hepes pH 7.5, 50 mM KCl, 10 mM Mg(CH_3_COO)_2_, 100 µg/mL CHX and cOmplete™ EDTA-free Protease Inhibitor Cocktail) prepared in a polypropylene centrifuge tube (14 × 95 mm; Beckman Coulter). A Gradient Master (Biocomp) was used to prepare the sucrose gradients. After centrifugation (285, 000 g for 2 h at 4 °C using a Beckman SW40Ti rotor), polysome profiles were recorded using an Äktaprime plus chromatography system (GE Healthcare). Proteins were precipitated with trichloroacetic acid (25 % (vol/vol); 15 min on ice). Following centrifugation at 18 000 g for 5 min at 4 °C, protein precipitates were washed with ice-cold acetone. After drying at room temperature, protein pellets were resuspended in 1x NuPAGE LDS Sample Buffer (Thermo Fisher Scientific). Vacant 80S monosomes were dissociated in 20 mM Hepes pH 7.5, 200 mM KCl, 10 mM Mg(CH_3_COO)_2_, 100 µg/mL CHX, cOmplete™ EDTA-free Protease Inhibitor Cocktail (Merck), 200 U/mL RNaseOUT, 0.5 % IGEPAL® CA-630 and 2 mM DTT.

### Purification of recombinant SBDS and EFL1 proteins

SBDS and EFL1 proteins were purified as previously described^13^.

### eIF6 release assay

#### Preparation of mature 80S ribosomes

Expanded mouse c-Kit+ bone marrow cells were lysed in ‘standard’ lysis buffer (20 mM Hepes pH 7.5, 50 mM KCl, 10 mM Mg(CH_3_COO)_2_, supplemented with cOmplete™ EDTA-free Protease Inhibitor Cocktail, 200 U/mL RNaseOUT inhibitor, 0.5 % IGEPAL® CA-630 and 2 mM DTT). A total of 150 A_260_ units of lysate was loaded on six sucrose gradients, and fractions corresponding to 80S monosomes were collected and further concentrated by centrifuging 30 min at 80000 g in a Beckman MLA-80 rotor fitted in an Optima MAX-XP ultracentrifuge. The sedimented 80S particles were resuspended in ‘standard’ buffer and aliquots stored at -80 °C.

#### Preparation of exogenous eIF6

c-Kit+ bone marrow cells isolated from transgenic eIF6 mice (genotype [*M2-rtTA/M2-rtTA*][*EIF6/+*]) were expanded and treated with doxycycline for 24 h to induce eIF6 overexpression. Cells were then harvested and lysed in ‘dissociation’ lysis buffer (20 mM Hepes pH 7.5, 500 mM KCl, 2 mM Mg(CH_3_COO)_2_ supplemented with 100 µg/mL CHX, cOmplete™ EDTA-free Protease Inhibitor Cocktail, 200 U/mL RNaseOUT inhibitor, 0.5 % IGEPAL® CA-630 and 2 mM DTT). A total of 29 A_260_ units of lysate was run on a single sucrose gradient, and the free fraction, which contains the vast majority of cellular eIF6 but is devoid of ribosomes, was collected and aliquots stored at -80 °C.

#### *In vitro* eIF6 release assay

In the first part of the assay, 10 uL (1.25 A_260_ units) of mature 80S particles were mixed with 100 uL of exogenous eIF6 in ‘dissociation’ buffer. The reaction mix was then incubated at 37 °C for 15 min both to promote the dissociation of the mature 80S particles into 40S and 60S subunits, and to allow the binding of the exogenous eIF6 to 60S subunits. The amount of eIF6 supplied was optimised empirically to be in slight excess over 60S subunits, thus saturating the available 60S subunits without a significant accumulation in the free fraction. In the second part of the protocol, the reaction mix was diluted with 500 uL of prewarmed KCl-free buffer (20 mM Hepes pH 7.5, 10 mM Mg(CH_3_COO)_2_), and incubated at 37 °C for 5 min to allow reassembly of 80S particles. Since the joining of 40S and 60S subunits into 80S particles is proportional to the release of eIF6 from 60S subunits, this experimental strategy allows the assessment of eIF6 release based on quantification of 80S to 60S ratio. The reaction mix was split equally into two tubes that were supplied either with 1 mM GTP or 1 mM GTP + 1250 nM SBDS + 600 nM EFL1. Following 1 h incubation at 25 °C, the reaction mixes were cooled down on ice, and loaded on sucrose gradients prepared in ‘standard’ buffer conditions.

### Electron cryo-microscopy sample preparation and data collection

c-Kit+ bone marrow cells were isolated from control mice that do not harbour *EIF6* transgene, and expanded keeping cell concentration below one million cells per mL. CHX-treated cells were lysed in ’standard’ lysis buffer as described in ’Polysome profiling experiments’. Following sucrose gradient sedimentation, 60S subunits from multiple gradients were pooled, sedimented by centrifugation (45 min at 80000 g in an MLA-80 rotor), resuspended in 20 mM HEPES pH 7.5, 50 mM KCl, 5 mM Mg(CH_3_COO)_2_ at a concentration of 100 mM, and stored at -80 °C. 60S ribosomal subunits were thawed on ice and centrifuged in a benchtop centrifuge for 10 minutes at 20000 g and the supernatant was carefully recovered. EM grids were prepared by depositing 3 µl of 60S subunits at 100 nM to freshly glow-discharged Quantifoil R 1.2/1.3 holey carbon grids (PELCO easyGlow). Grids were then blotted with a Vitrobot Mark IV (FEI company) using the following parameters: blot time 1 s, blot force -7, wait time 10 s, no drain time. Blotted grids were finally vitrified in liquid ethane and stored in liquid nitrogen. Grids were screened on a Tecnai T12 microscope (FEI Company) and data acquisition performed under low-dose conditions on a Titan Krios microscope (FEI Company) operated at 300 kV over 24 h. The dataset was recorded on a Falcon III detector (FEI Company) at a nominal magnification of 75,000x (effective pixel size of 1.10 Å on the object scale) with a defocus range of −0.8 to −3.2 µm and a total dose of ∼77 e−/Å^2^ accumulated over 2 s exposures in 38 fractions. The acquisition of 3024 movies was performed semi-automatically using EPU software (FEI Company).

### Electron cryo-microscopy data processing

Data processing was handled within the RELION software package^48–50^. Movies were first corrected for motion using Motioncor2^51^ and CTF was estimated by CTFFIND4^52^. Particles were then picked using the Laplacian-of-Gaussian (LoG) filter, extracted and 2D classified in RELION. 3D classification was then performed on selected 2D classes to further discard non-ribosomal and contaminating 80S particles. The 60S ribosomal subunit-containing class was selected for 3D auto-refinement to generate a consensus map. Masking and auto-sharpening was done through post-processing in RELION to obtain the final high-resolution map.

To quantify the proportion of eIF6-bound ribosomal particles, we made use of a combination of particle subtraction and 3D masked classifications in RELION (Figure 1B). We first focused on the L1-stalk to sort particles relative to their maturation state. We generated a mask around the L1-stalk and the tRNA E-site from the consensus map and used it in 3D masked classification, leading to the isolation of mature ribosomal particles (88% of consensus-refined particles). We then generated a soft-edged mask around the area of the eIF6 binding site from the consensus map as an input. Signal outside this mask was subtracted in the newly obtained mature ribosomal particles subset. We finally generated 3D classes focusing on the area inside the mask. 4 classes were obtained, of which 3 showed clear density inside the masked area indicating the unequivocal presence of eIF6 and were then pooled for quantification (83% of mature ribosomal particles).

### Immunoblotting

Proteins in 1x NuPAGE LDS sample buffer (with 50 mM DTT) were incubated at 80 °C for 10 min and run on NuPAGE Bis-Tris polyacrylamide gels in NuPAGE MOPS SDS running buffer (Thermo Fisher Scientific). The iBlot 2 gel transfer device (Thermo Fisher Scientific) was used to transfer proteins to nitrocellulose membranes. Membranes were blocked in 5 % milk in PBS supplemented with 0.1% tween (NBS Biologicals) for 1 h, and subsequently incubated with appropriate primary antibodies overnight at 4 °C on a shaker. After 3x 10 min washes with PBS-tween, the blots were incubated with the appropriate secondary horseradish peroxidase-conjugated antibody at room temperature for 1 h followed by detection using the SuperSignal West Pico PLUS reagents (Thermo Fisher Scientific). For a full list of antibodies, see Supplementary Table 2.

### Subcellular fractionation

1 × 10^6^ freshly isolated bone marrow cells from Dox-treated mice were washed twice with ice-cold PBS, and resuspended in 0.5 mL of ‘standard’ lysis buffer (20 mM Hepes pH 7.5, 50 mM KCl, 10 mM Mg(CH_3_COO)_2_, cOmplete™ EDTA-free Protease Inhibitor Cocktail, 0.5 % IGEPAL® CA-630, and 2 mM DTT. Following 30 min incubation on ice, lysates were centrifuged for 5 min at 7000 g to pellet the nuclei, while supernatants representing the cytosolic fraction were collected. Following a wash with ice-cold PBS and centrifugation as above, the nuclear pellet was resuspended in 0.5 mL of ice-cold RIPA buffer supplied with 1 U/mL Benzonase (Merck), and incubated on a rotator for 1 h at 4 °C. Following centrifugation for 5 min at 7000 g to pellet insoluble material, supernatant representing the nuclear fraction was collected. Finally, both fractions were resuspended in 1x NuPAGE LDS sample buffer with 50 mM DTT.

### Quantitative real-time PCR

Total RNA was isolated from FACS-purified cells using the RNeasy mini kit (Qiagen). cDNA was transcribed with SuperScript III reverse trancriptase (Thermo Fisher Scientific). Real-time PCR reactions were performed using the SsoFast™ EvaGreen^®^ Supermix (Bio-Rad) and ABI 7900HT fast Real-time PCR system (Thermo Fisher Scientific). Primers are listed in Supplementary Table 3.

### Light microscopy

*Drosophila* was maintained using standard culture techniques. All crosses were performed at 25 °C. Fly strains and genotypes are described in Supplementary Tables 4 **and** 5. Whole *Drosophila* samples were collected at 1, 3, 5, and 11 days after egg laying (AEL). Larvae were fixed with 4% paraformaldehyde and adult flies were frozen before photography. For the imaging of fly eyes, two to four day old *Drosophila* adults were frozen at -20 °C for one hour. Both whole fly and adult eye photographs were collected using a Nikon SMZ18 microscope with NIS-Elements D (version 4.40).

### Scanning Electron Microscopy

*Drosophila* adult eye samples were prepared as described^53^. Samples were viewed on a Philips XL30 scanning electron microscope.

### Immunostaining

*Drosophila* wing discs dissected from third instar larvae in culture medium (*Drosophila* M3 media (Sigma), 10 % FCS (Sigma) and P/S (Sigma) were collected within 10 min into culture medium containing 50 µM of OP-Puro (Invitrogen) and kept in a 25 °C incubator for 30 min. Wing discs were then washed twice with ice-cold PBS (Invitrogen) with 1% BSA (Sigma) and 100 µg/ml CHX (Sigma). Wing discs were fixed and permeabilised using the Cytofix/Cytoperm Fixation Permeabilization Kit (BD Biosciences). Azide-alkyne cycloaddition was performed using the Click-iT Cell Reaction Buffer Kit (Invitrogen) with azide conjugated to Alexa Fluor 596 at 5 µM final concentration. Following a 30 min reaction, wing discs were washed three times in PBS and mounted on slides in medium containing DAPI (Vector). Images were collected on a Zeiss LSM710 confocal system and imported to Image J 10.4 (Image J) and Photoshop (Adobe 2020), and adjusted for brightness and contrast uniformly across entire fields.

## ACKNOWLEDGEMENTS

This work was supported by the Swedish Childhood Cancer Fund (PDS13/001, to PJ), the Swedish Research Council (2014-06807, to PJ), a Specialist Programme from Bloodwise (12048, to AJW), the UK Medical Research Council (MC_U105161083, to AJW), a Wellcome Trust strategic award to the Cambridge Institute for Medical Research (100140, to AJW), a core support grant from the Wellcome Trust and MRC to the Wellcome Trust-Medical Research Council Cambridge Stem Cell Institute, the Connor Wright Project, Ted’s Gang and the Cambridge National Institute for Health Research Biomedical Research Centre. C.C.W. acknowledges funding from a Wellcome Intermediate Fellowship (Grant Number: 105914/Z/14/Z) and the Kay Kendall Leukaemia Fund. We thank Dr. D. Y. Chirgadze for assistance with data collection at the Cryo-EM Facility, Department of Biochemistry, University of Cambridge, funded by the Wellcome Trust (206171/Z/17/Z; 202905/Z/16/Z), the Departments of Biochemistry and Chemistry, the Schools of Biological Sciences and Clinical Medicine and the University of Cambridge. We thank R. Grenfell at the Cancer Research UK Cambridge Institute for assistance with imaging flow cytometry. The Cambridge NIHR BRC Cell Phenotyping Hub supported this research. We thank T. Hamilton, D. Pask, D. Kent, G. Giotopoulos, B. Huntly and A.R. Green for facilitating animal experiments. We thank M. Freeman, the Bloomington *Drosophila* Stock Center, NIG-Fly Stock Center and Vienna *Drosophila* RNAi Center for providing fly stocks.

## AUTHORSHIP

Conflict-of-interest disclosure: The authors declare no competing financial interests.

---

**Supplementary Figure 1.**
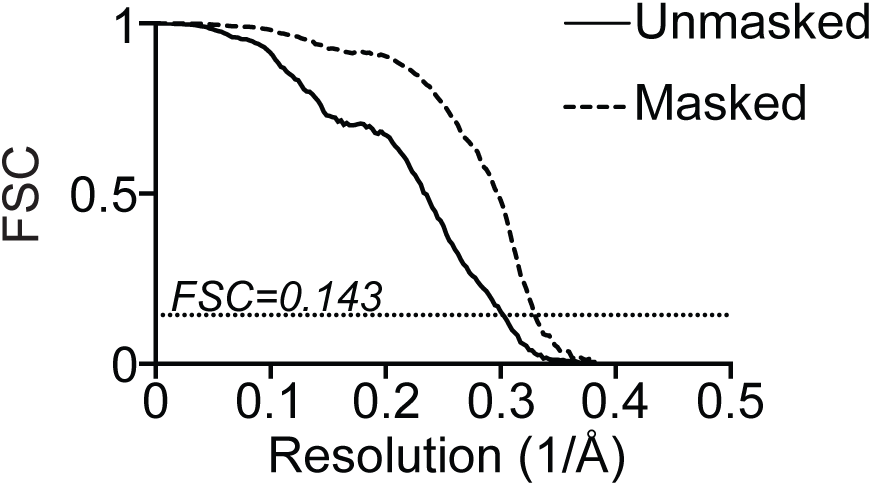
Fourier shell correlation curves of the cryo-EM data set. Fourier shell correlation curves of both solvent masked and unmasked final maps indicating maximum resolution at 0.143 ‘gold standard’ threshold.

**Supplementary Figure 2.**
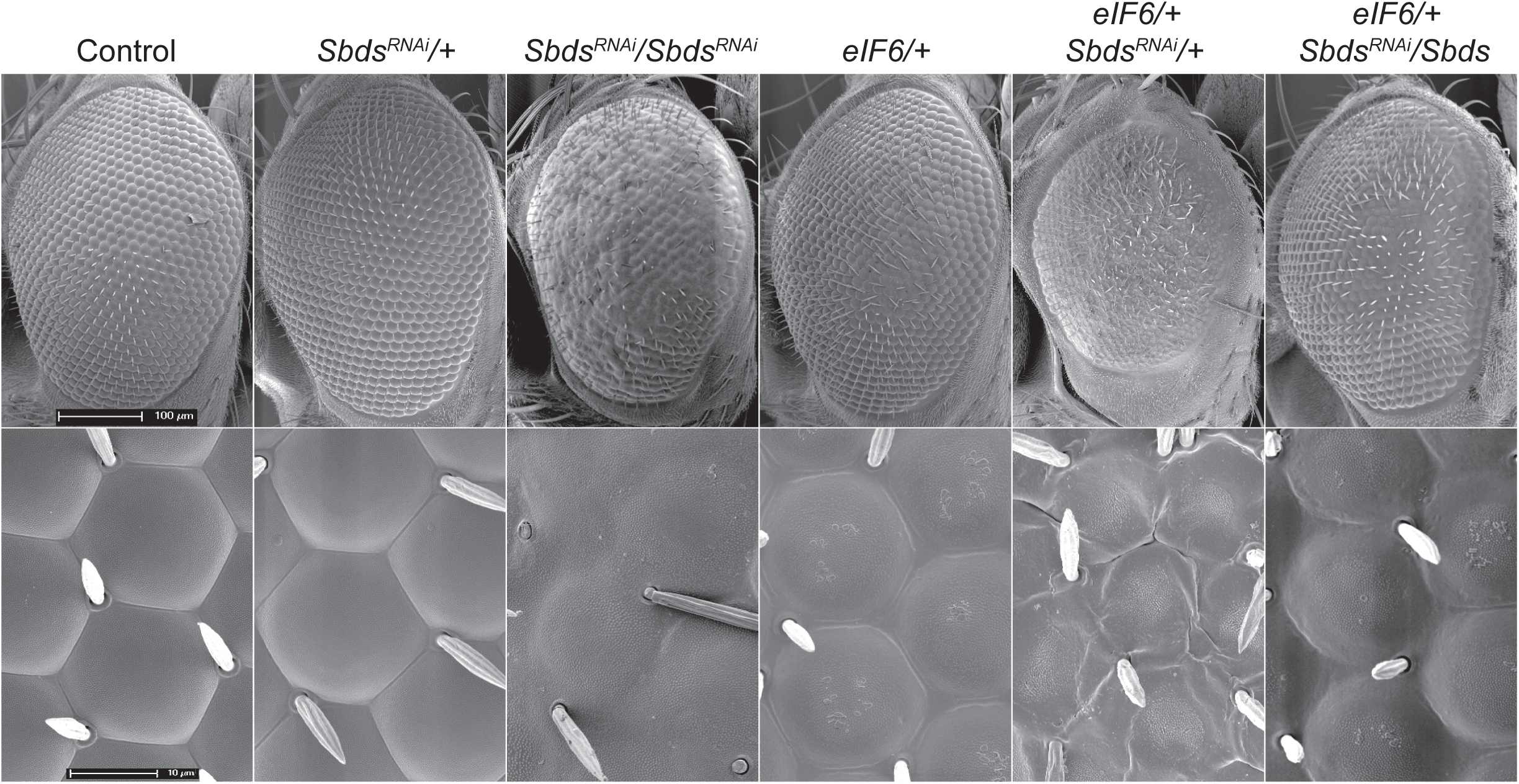
Genetic interaction between eIF6 and Sbds. Representative scanning electron microscopy images showing the *Drosophila* eye phenotypes in the indicated genotypes. n=3.

**Supplementary Figure 3.**
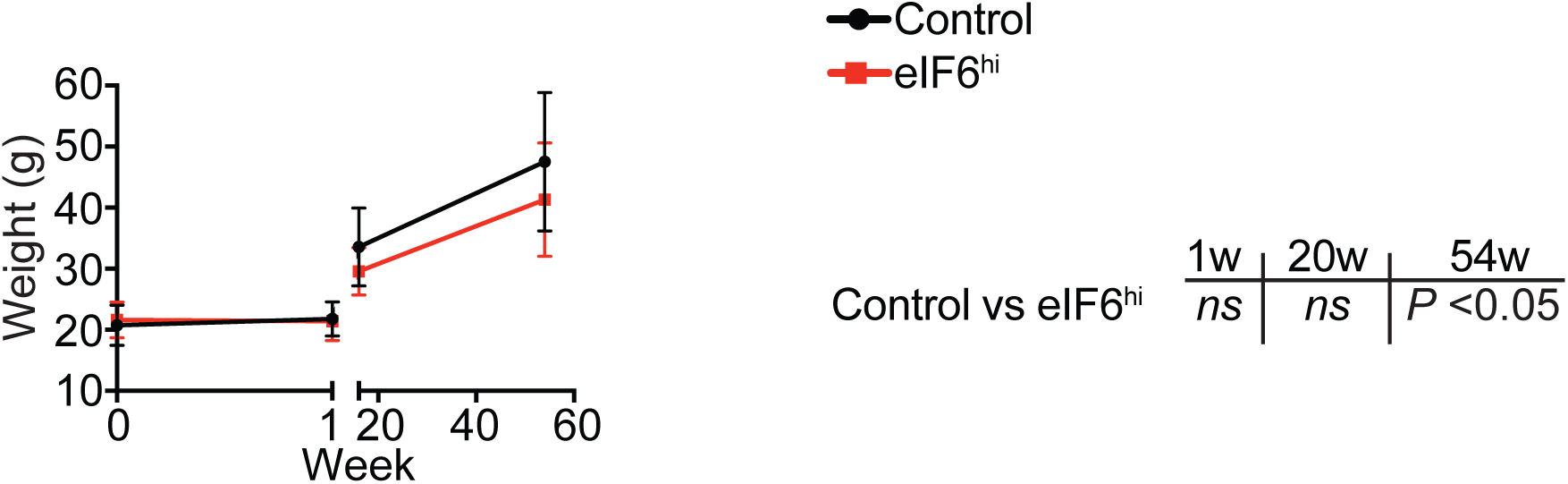
Dox administration has no acute impact on weight gain in eIF6^hi^ mice. Time represents weeks fed on doxycycline diet. n=9-12 per genotype. Student’s *t* test was used to determine statistical significance, and two-tailed *P* values are shown. Error bars represent standard deviation.

**Supplementary Figure 4.**
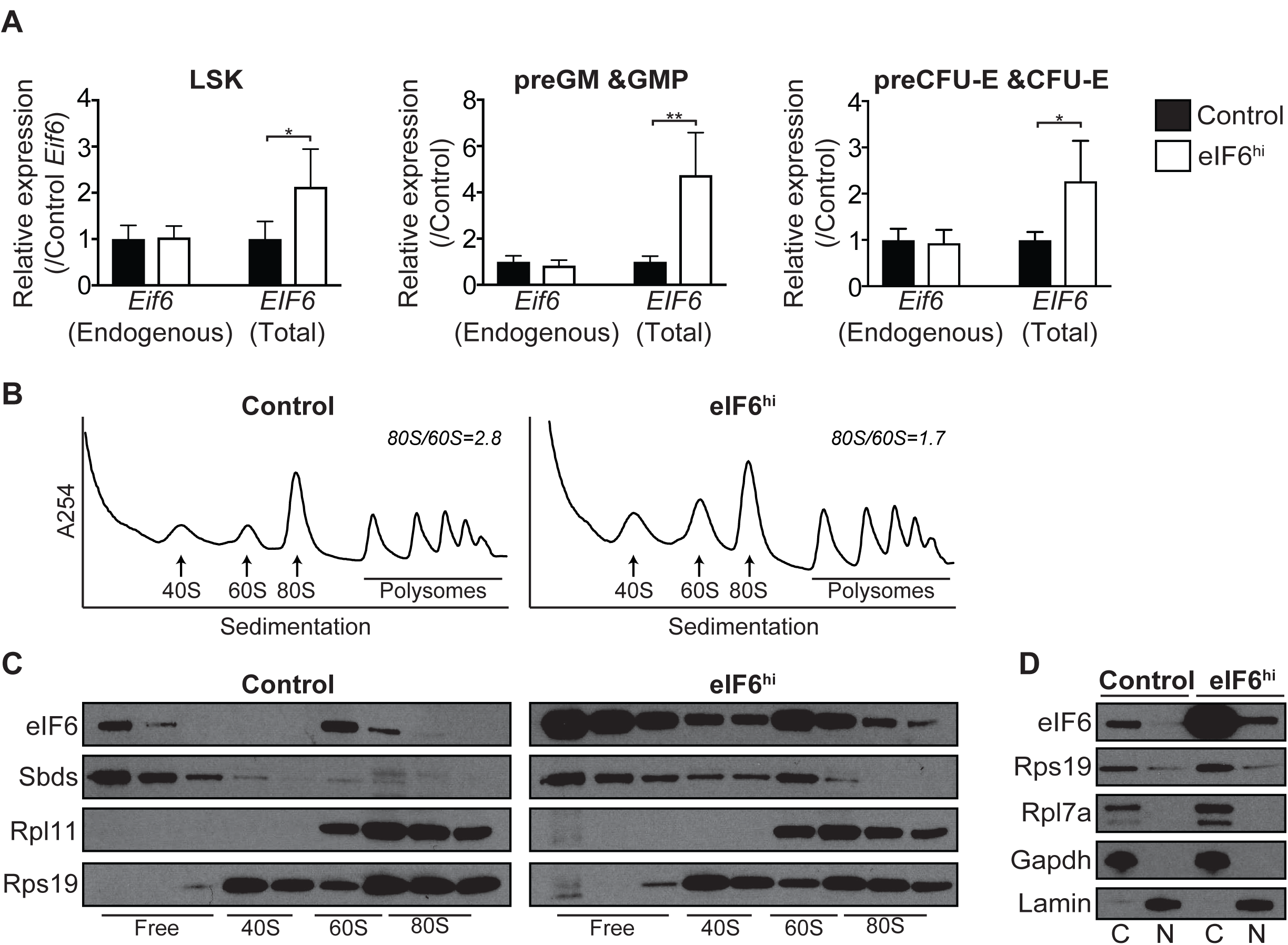
Validation of *EIF6* transgene expression in eIF6^hi^ mice. (A) Quantitative real-time PCR analysis of *EIF6* mRNA expression in LSK, preGM/GMP and preCFU-E/CFU-E progenitor cells isolated from adult mice after two weeks of doxycycline administration. (n= 4 per genotype). (B) Sucrose gradient sedimentation of extracts from cultured c-Kit+ bone marrow cells derived from the indicated mouse strains. Dox induction, 24 h. (C) Immunoblotting analysis to visualise eIF6, Sbds, Rpl11 and Rps19 across the sucrose density gradients shown in (B). (D) Subcellular fractionation of freshly isolated unfractionated bone marrow cells from adult mice after two weeks of doxycycline administration. C= cytoplasmic fraction, N= nuclear fraction. LSK=Lineage- Sca-1+ c-Kit+, preGM= pre-granulocyte-macrophage progenitor, GMP= granulocyte-macrophage progenitor, pre-CFU-E= pre-colony-forming unit-erythroid progenitor. Student’s *t* test was used to determine statistical significance, and two-tailed *P* values are shown. Error bars represent standard deviation.

**Supplementary Figure 5.**
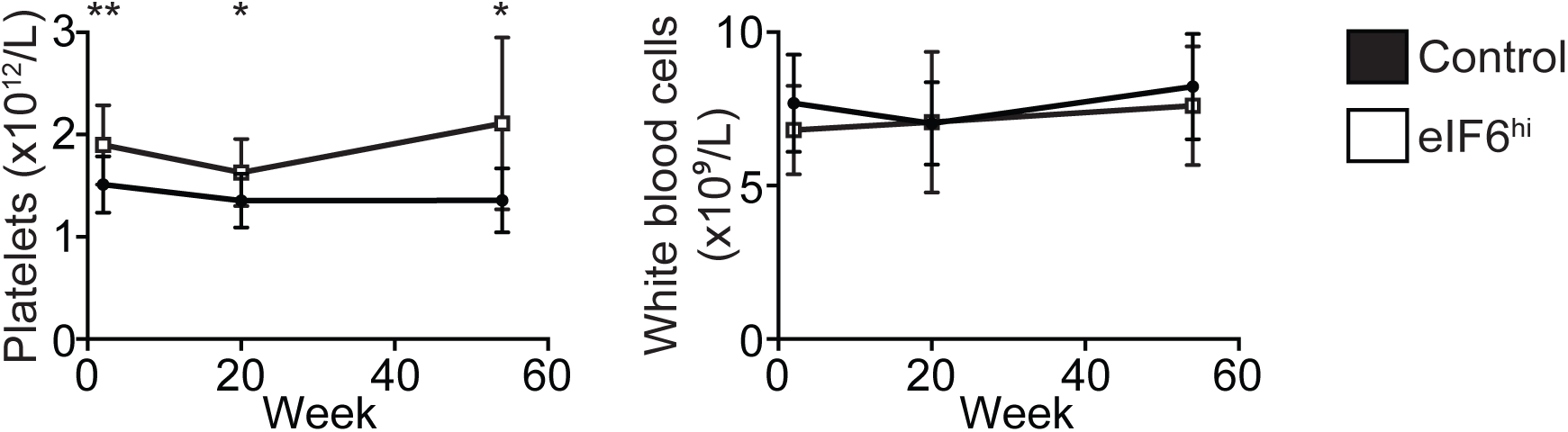
Peripheral blood analysis of eIF6^hi^ mice after two weeks of doxycycline induction. n=12-15 per genotype. Student’s *t* test was used to determine statistical significance and two-tailed *P* values are shown. Error bars represent standard deviation.

**Supplementary Figure 6.**
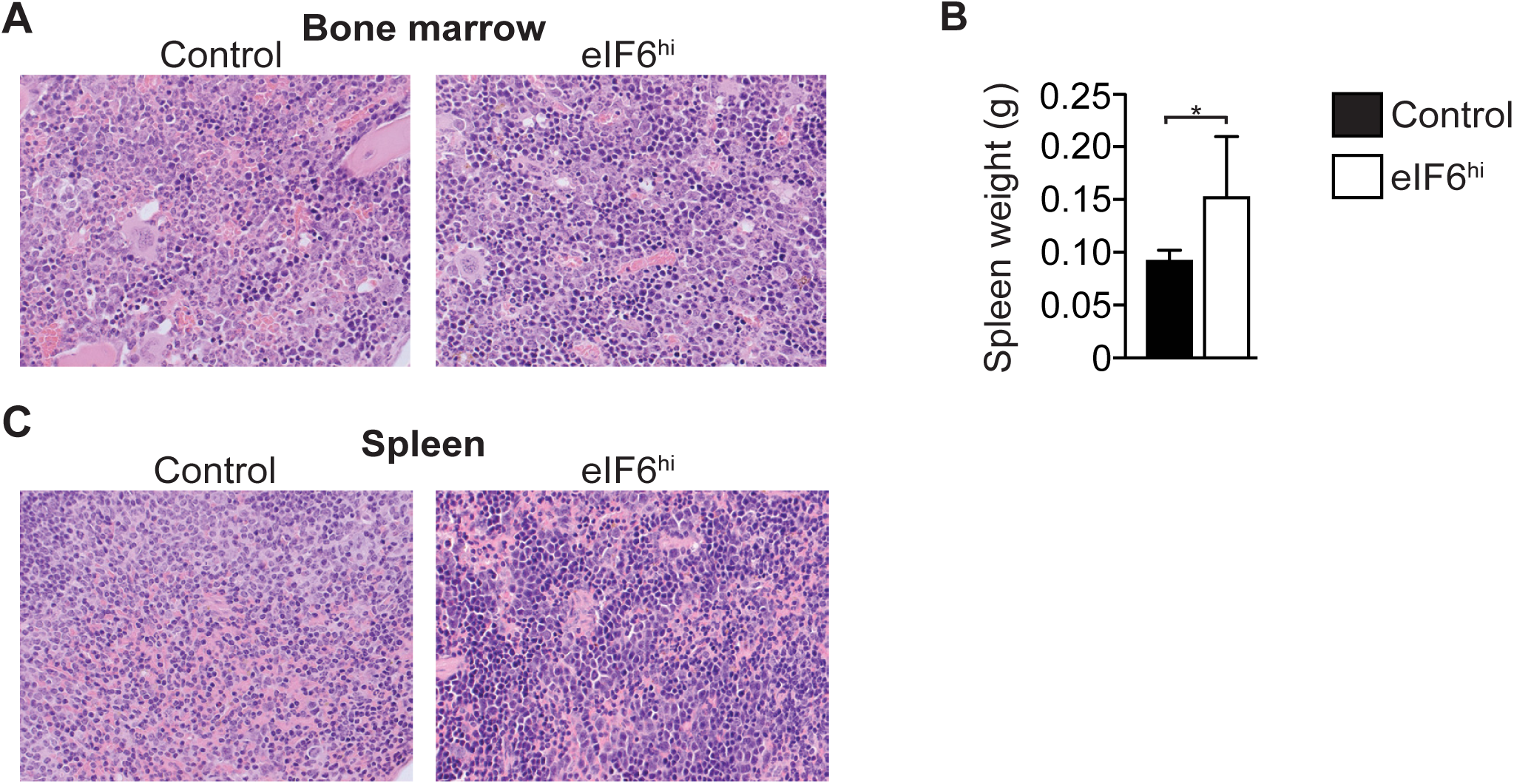
Histological examination of the eIF6^hi^ mice. (A) Representative bone marrow sections (40X). (B) Spleen weight (n=7 per genotype). (C) Representative spleen sections (40X). Student’s *t* test was used to determine statistical significance, and two-tailed *P* values are shown. Error bars represent standard deviation.

**Supplementary Figure 7.**
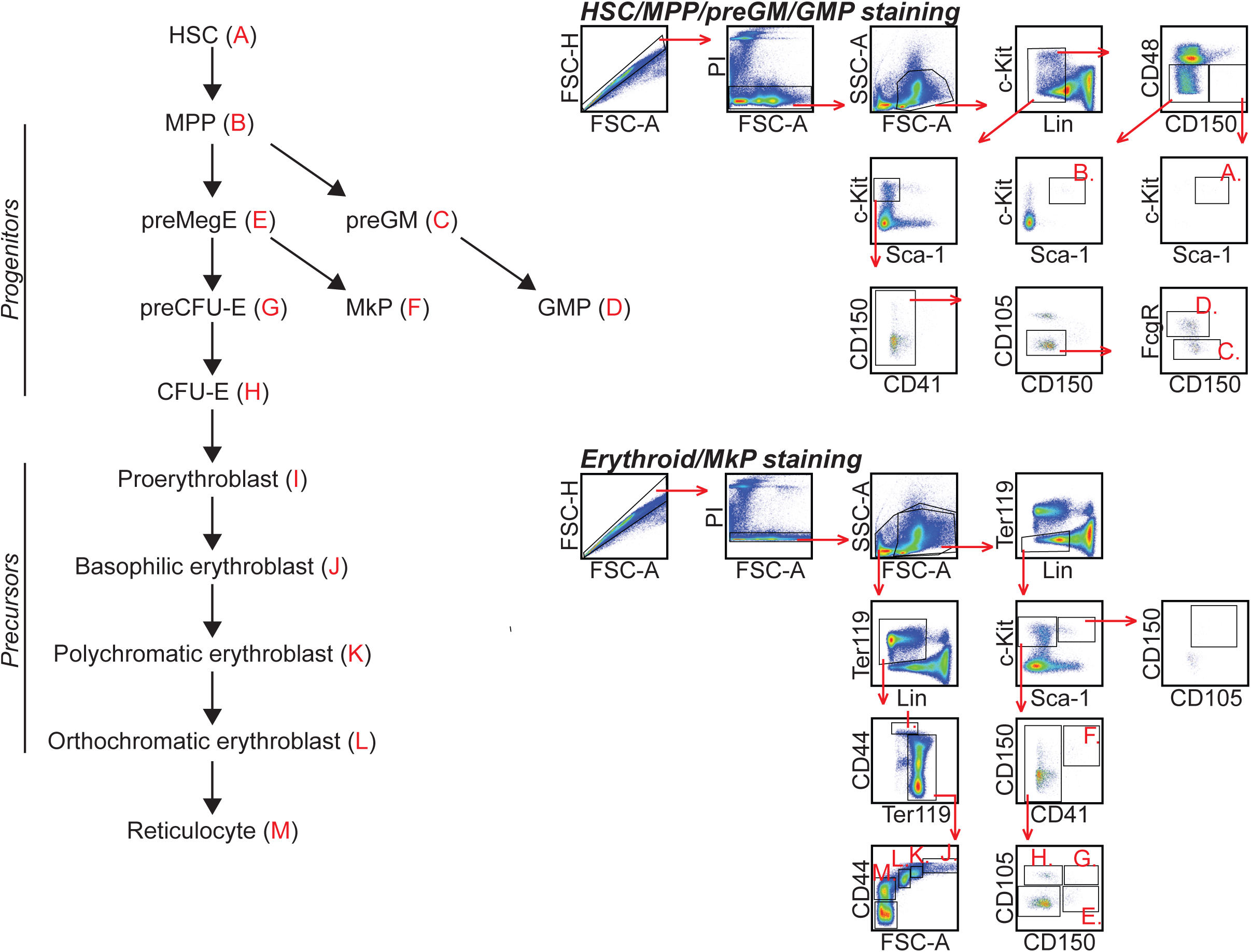
Flow cytometry strategy to quantify bone marrow subpopulations.

**Supplementary Figure 8.**
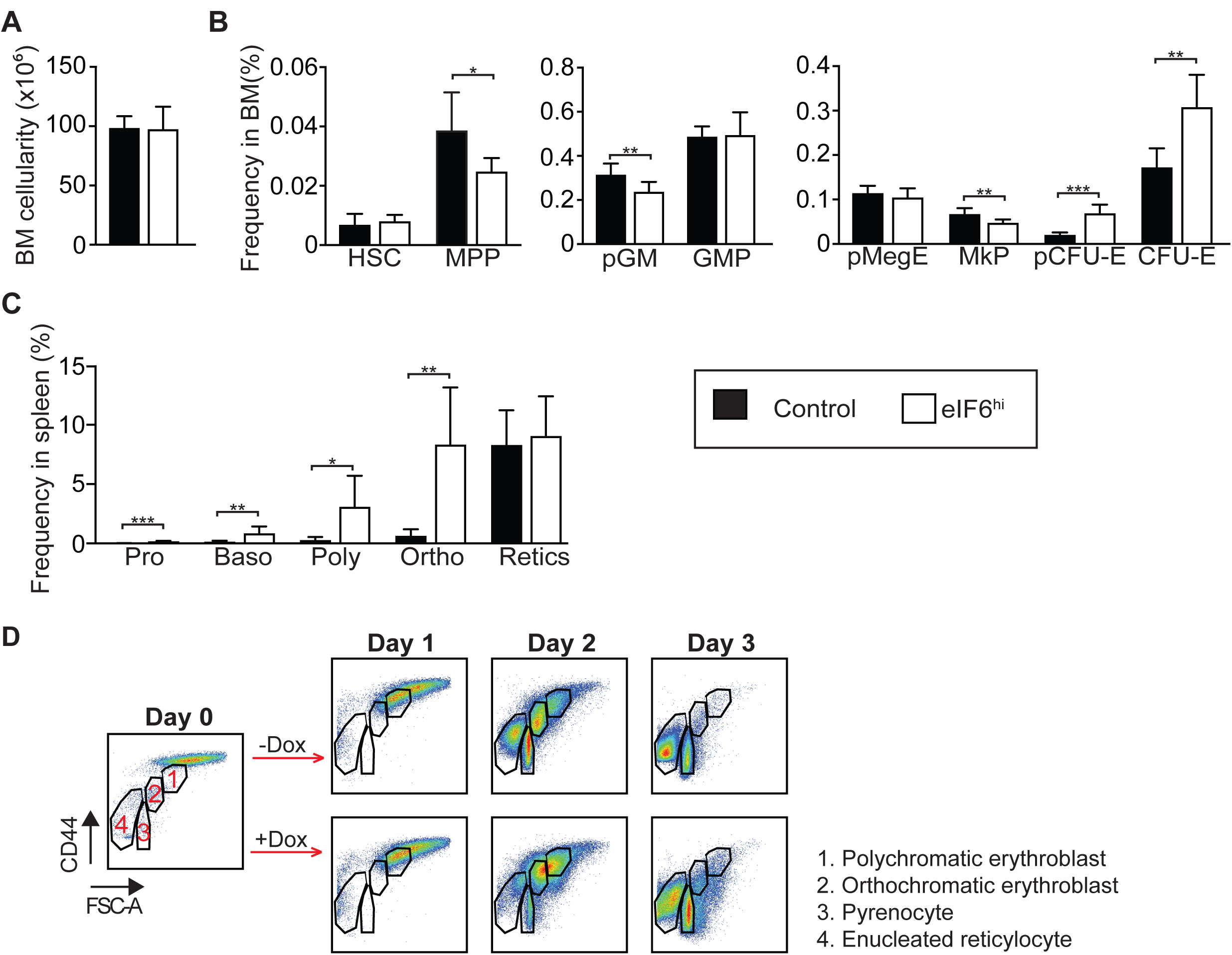
Flow cytometric analysis of the haematopoietic phenotype of the eIF6^hi^ mice. (A) Bone marrow cellularity and (B) frequency of haematopoietic stem and progenitor cells in the bone marrow after two weeks of Dox administration (n=7 per genotype). (C) Frequency of erythroid precursor cells in the spleen after two weeks of Dox administration (n=7 per genotype). (D) Representative *in vitro* differentiation culture of prospectively purified CFU-E/proerythroblast cells (n=2). Fresh cells were isolated from the bone marrow of Dox-treated eIF6^hi^ mice, and depleted for TER-119, Gr-1, CD11b, CD4, CD8, B220, CD41, CD16/32, CD150 and Sca-1. Enriched cells were cultured with or without Dox. HSC = haematopoietic stem cell; MPP = multipotent progenitor; pGM= pre-granulocyte-macrophage progenitor; GMP = granulocyte-macrophage progenitor, preMegE = pre-megakaryocyte-erythroid progenitor, MkP = megakaryocyte progenitor, preCFU-E = pre-colony-forming unit-erythroid progenitor. Student’s *t* test was used to determine statistical significance, and two-tailed *P* values are shown. Error bars represent standard deviation.

**Supplementary Figure 9.**
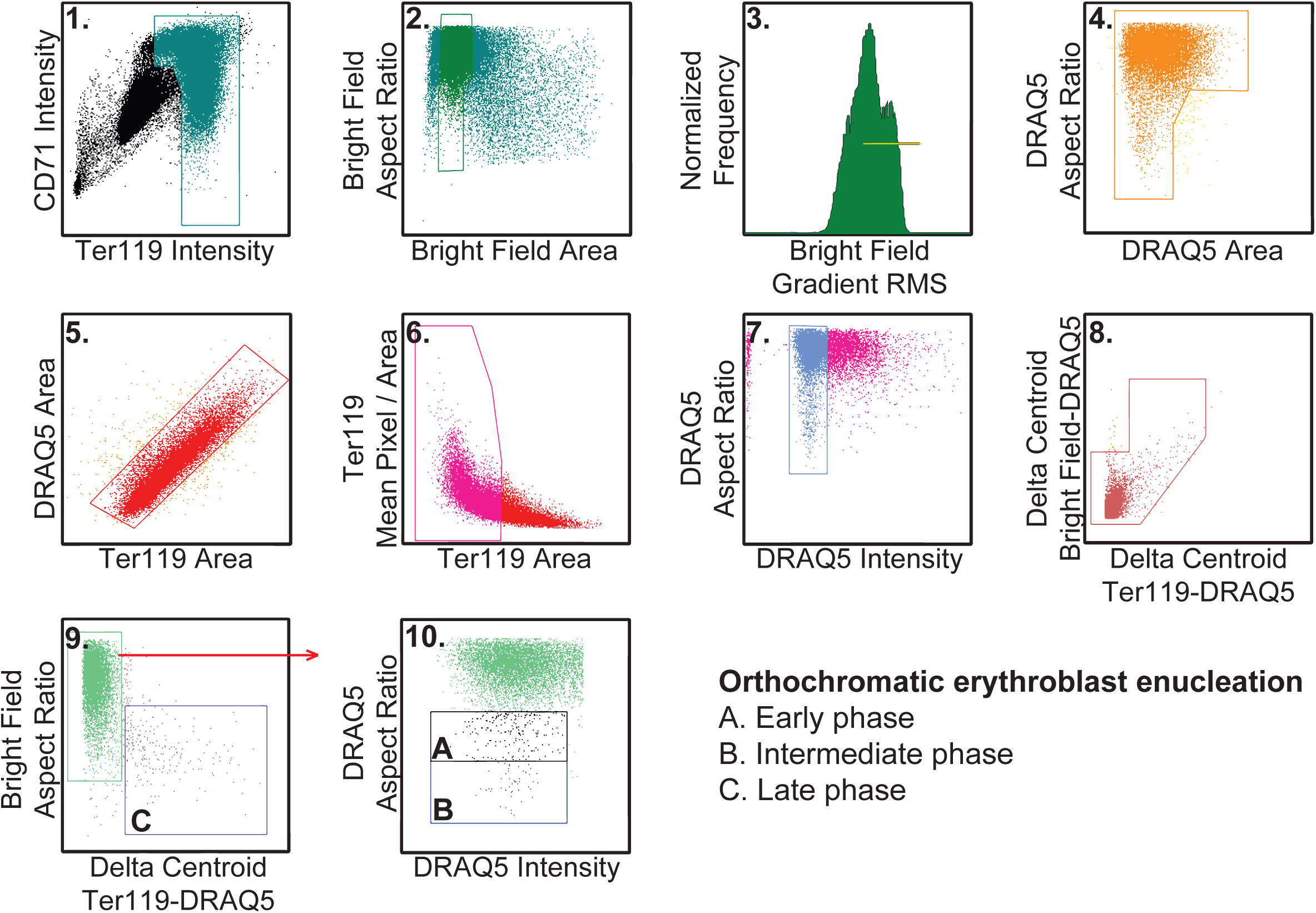
IDEAS strategy to visualise enucleating erythroblasts in the bone marrow.

**Supplementary Figure 10.**
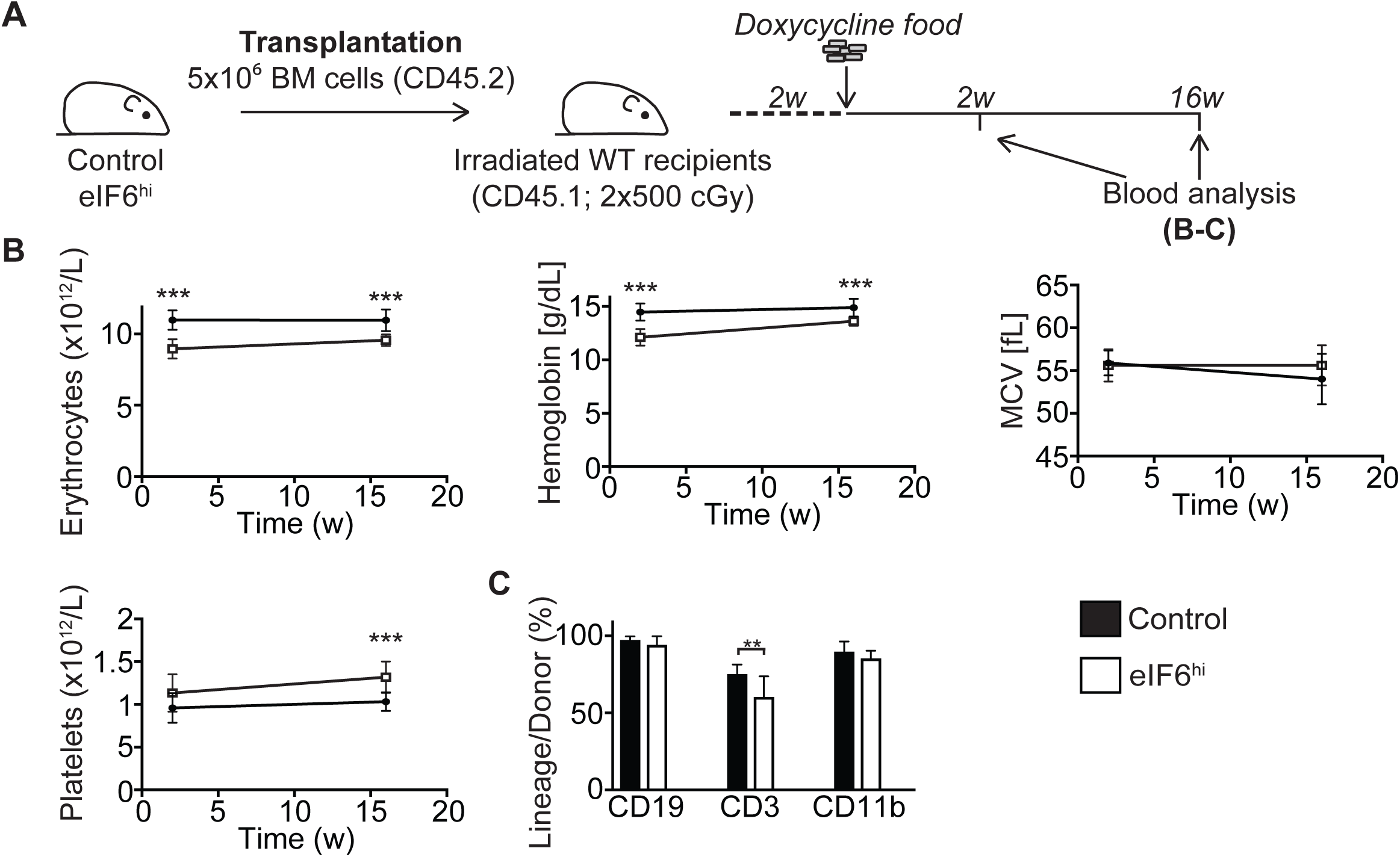
The haematopoietic phenotype in eIF6^hi^ mice is autonomous to the blood system. (A) Overview of the transplantation strategy. Five million freshly isolated unfractionated bone marrow cells from uninduced control or eIF6^hi^ mice were transplanted into the tail vein of lethally irradiated (2x 500 cGy) wild-type recipients (CD45.1). Two weeks after transplantation, Dox was administered to the recipient mice to induce expression of the *EIF6* transgene, and peripheral blood cellularity was analysed at indicated time-points. (B) The number of erythrocytes and platelets, hemoglobin concentration and mean corpuscular volume (MCV) in the peripheral blood over time (n= 9-10 per group). (C) Donor cell reconstitution within the different white blood cell lineages after 16 weeks of Dox administration (n=9-10 per group). Student’s *t* test was used to determine statistical significance, and two-tailed *P* values are shown. Error bars represent standard deviation.

**Supplementary Table 1.**
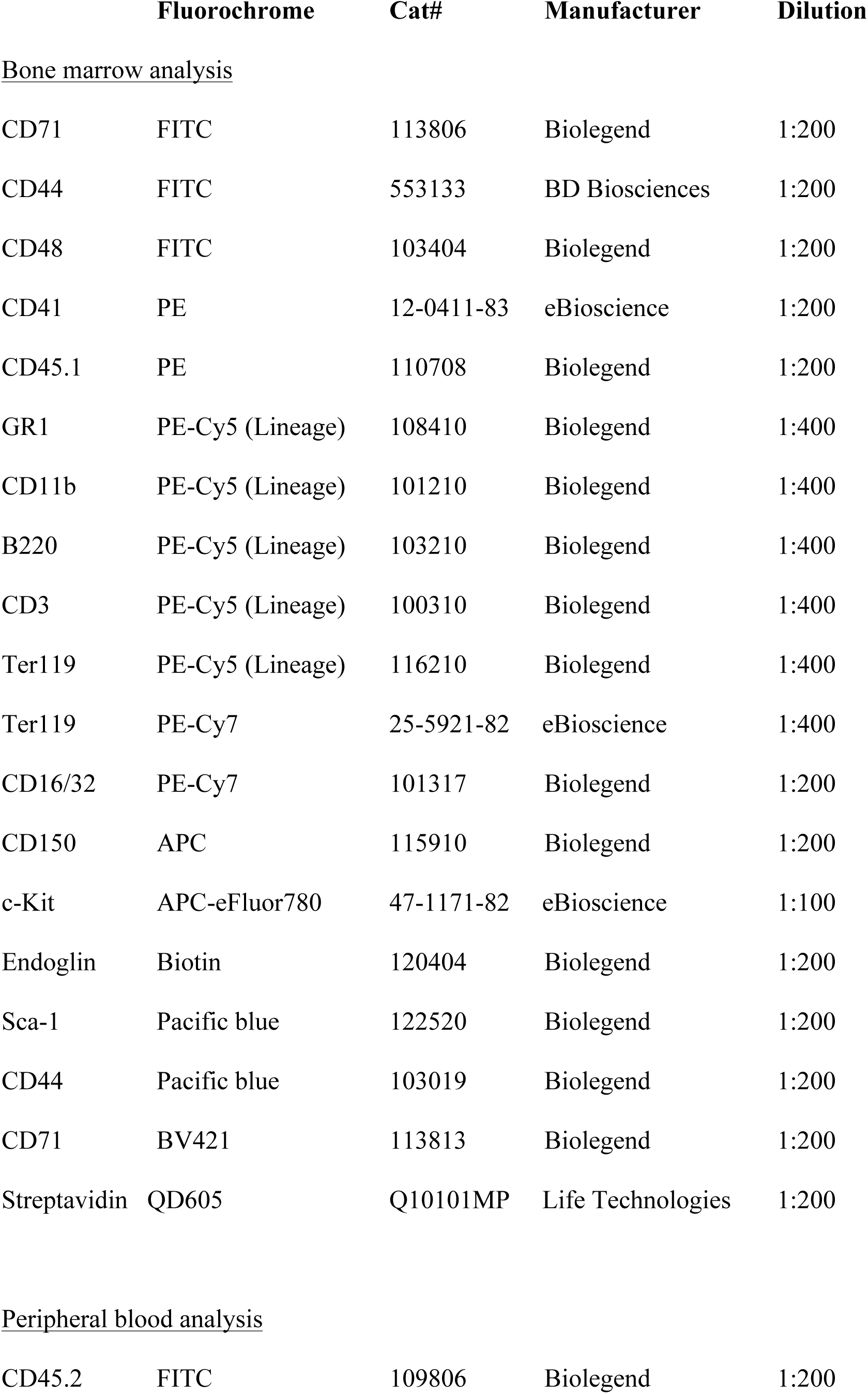
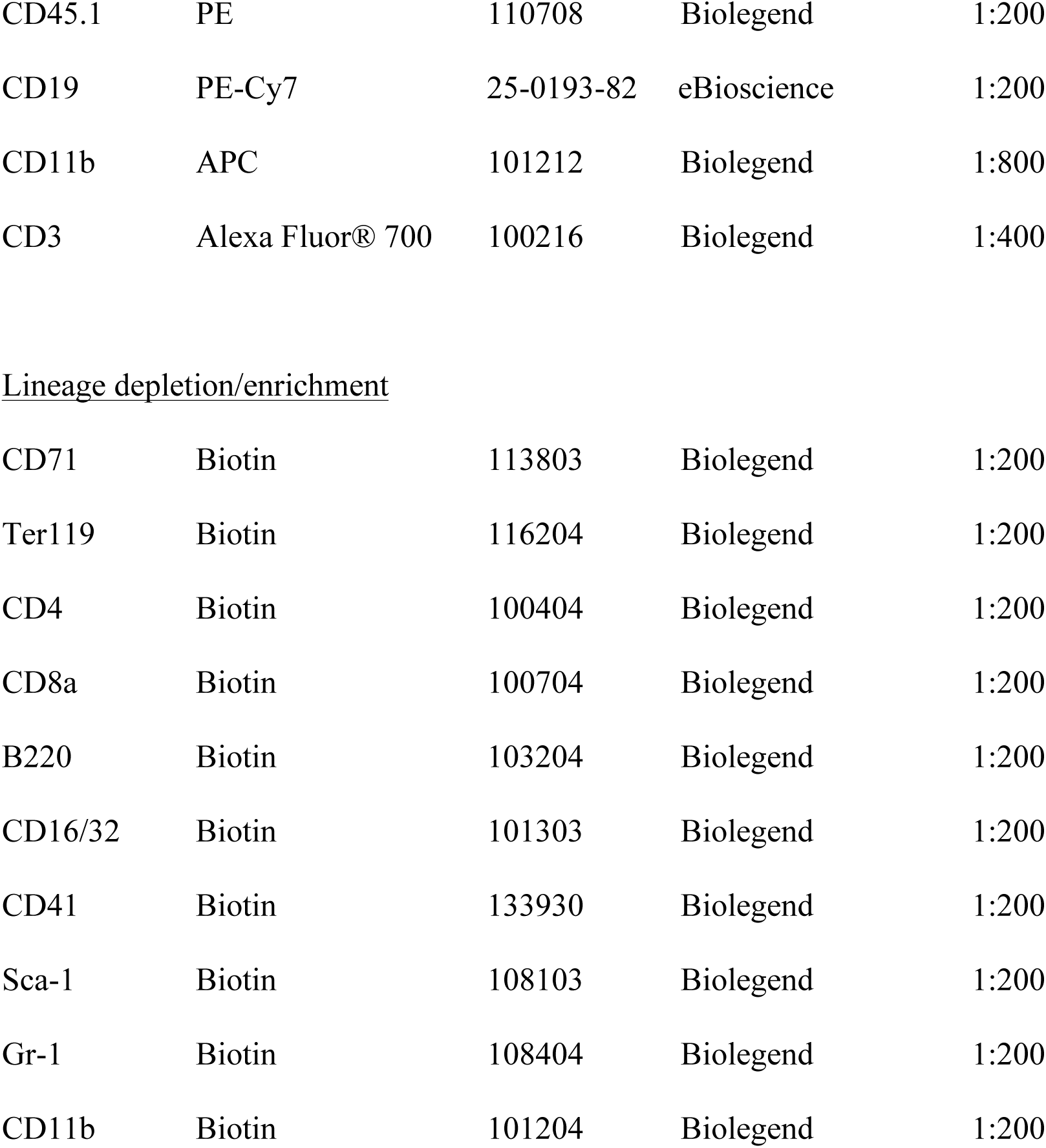
Antibodies and reagents used in flow cytometry.

|  | **Fluorochrome** | **Cat#** | **Manufacturer** | **Dilution** |
| --- | --- | --- | --- | --- |
| <u>Bone marrow analysis</u> |  |  |  |  |
| CD71 | FITC | 113806 | Biolegend | 1:200 |
| CD44 | FITC | 553133 | BD Biosciences | 1:200 |
| CD48 | FITC | 103404 | Biolegend | 1:200 |
| CD41 | PE | 12-0411-83 | eBioscience | 1:200 |
| CD45.1 | PE | 110708 | Biolegend | 1:200 |
| GR1 | PE-Cy5 (Lineage) | 108410 | Biolegend | 1:400 |
| CD11b | PE-Cy5 (Lineage) | 101210 | Biolegend | 1:400 |
| B220 | PE-Cy5 (Lineage) | 103210 | Biolegend | 1:400 |
| CD3 | PE-Cy5 (Lineage) | 100310 | Biolegend | 1:400 |
| Ter119 | PE-Cy5 (Lineage) | 116210 | Biolegend | 1:400 |
| Ter119 | PE-Cy7 | 25-5921-82 | eBioscience | 1:400 |
| CD16/32 | PE-Cy7 | 101317 | Biolegend | 1:200 |
| CD150 | APC | 115910 | Biolegend | 1:200 |
| c-Kit | APC-eFluor780 | 47-1171-82 | eBioscience | 1:100 |
| Endoglin | Biotin | 120404 | Biolegend | 1:200 |
| Sca-1 | Pacific blue | 122520 | Biolegend | 1:200 |
| CD44 | Pacific blue | 103019 | Biolegend | 1:200 |
| CD71 | BV421 | 113813 | Biolegend | 1:200 |
| Streptavidin | QD605 | Q10101MP | Life Technologies | 1:200 |
| <u>Peripheral blood analysis</u> |  |  |  |  |
| CD45.2 | FITC | 109806 | Biolegend | 1:200 |
| CD45.1 | PE | 110708 | Biolegend | 1:200 |
| CD19 | PE-Cy7 | 25-0193-82 | eBioscience | 1:200 |
| CD11b | APC | 101212 | Biolegend | 1:800 |
| CD3 | Alexa Fluor® 700 | 100216 | Biolegend | 1:400 |
| Lineage depletion/enrichment |  |  |  |  |
| CD71 | Biotin | 113803 | Biolegend | 1:200 |
| Ter119 | Biotin | 116204 | Biolegend | 1:200 |
| CD4 | Biotin | 100404 | Biolegend | 1:200 |
| CD8a | Biotin | 100704 | Biolegend | 1:200 |
| B220 | Biotin | 103204 | Biolegend | 1:200 |
| CD16/32 | Biotin | 101303 | Biolegend | 1:200 |
| CD41 | Biotin | 133930 | Biolegend | 1:200 |
| Sca-1 | Biotin | 108103 | Biolegend | 1:200 |
| Gr-1 | Biotin | 108404 | Biolegend | 1:200 |
| CD11b | Biotin | 101204 | Biolegend | 1:200 |

**Supplementary Table 2.**
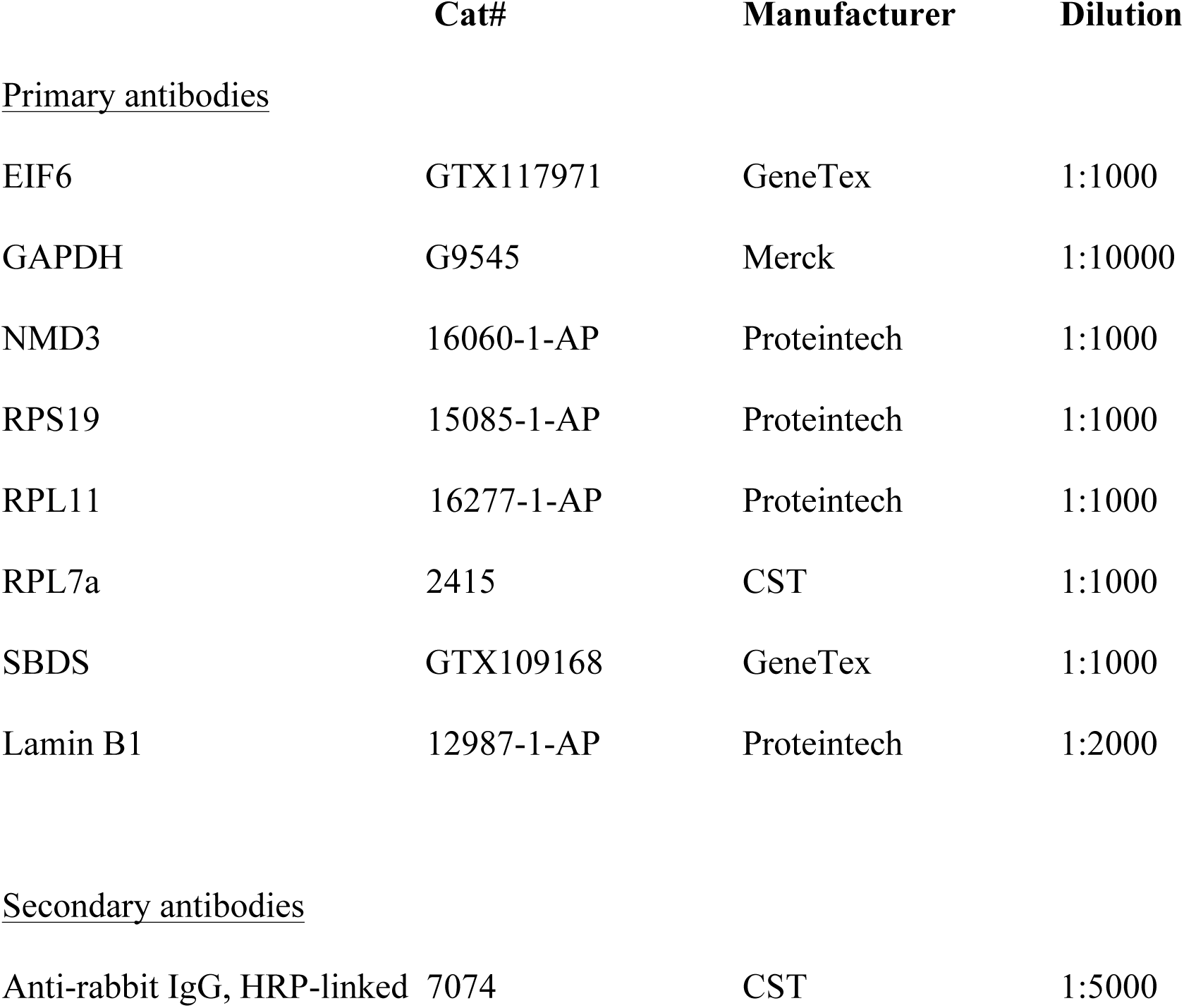
Antibodies used in immunoblotting.

|  | **Cat#** | **Manufacturer** | **Dilution** |
| --- | --- | --- | --- |
| <u>Primary antibodies</u> |  |  |  |
| EIF6 | GTX117971 | GeneTex | 1:1000 |
| GAPDH | G9545 | Merck | 1:10000 |
| NMD3 | 16060-1-AP | Proteintech | 1:1000 |
| RPS19 | 15085-1-AP | Proteintech | 1:1000 |
| RPL11 | 16277-1-AP | Proteintech | 1:1000 |
| RPL7a | 2415 | CST | 1:1000 |
| SBDS | GTX109168 | GeneTex | 1:1000 |
| Lamin B1 | 12987-1-AP | Proteintech | 1:2000 |
| <u>Secondary antibodies</u> |  |  |  |
| Anti-rabbit IgG, HRP-linked | 7074 | CST | 1:5000 |

**Supplementary Table 3.**
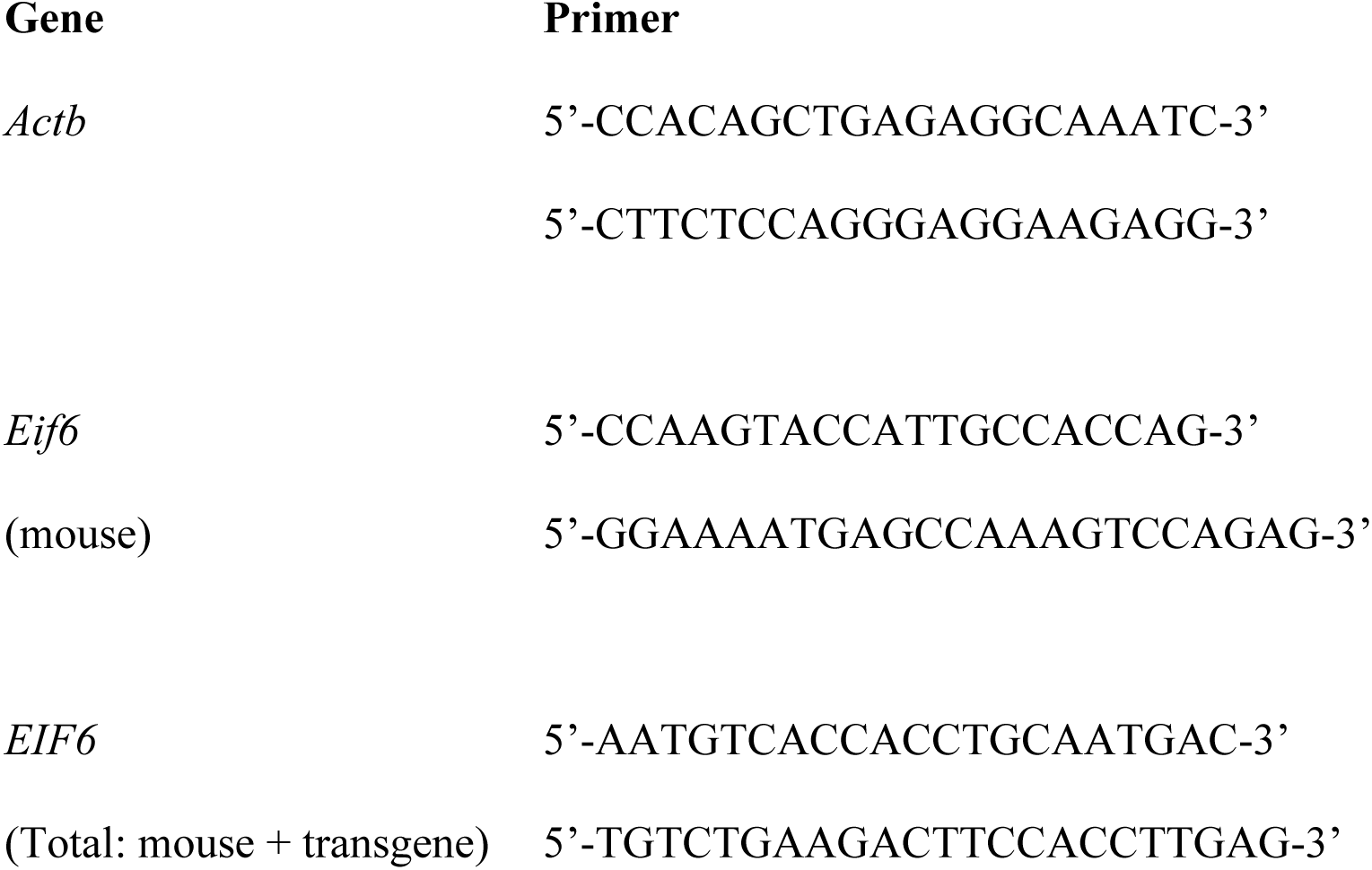
Custom primers used for quantitative real-time PCR.

**Supplementary Table 4.**
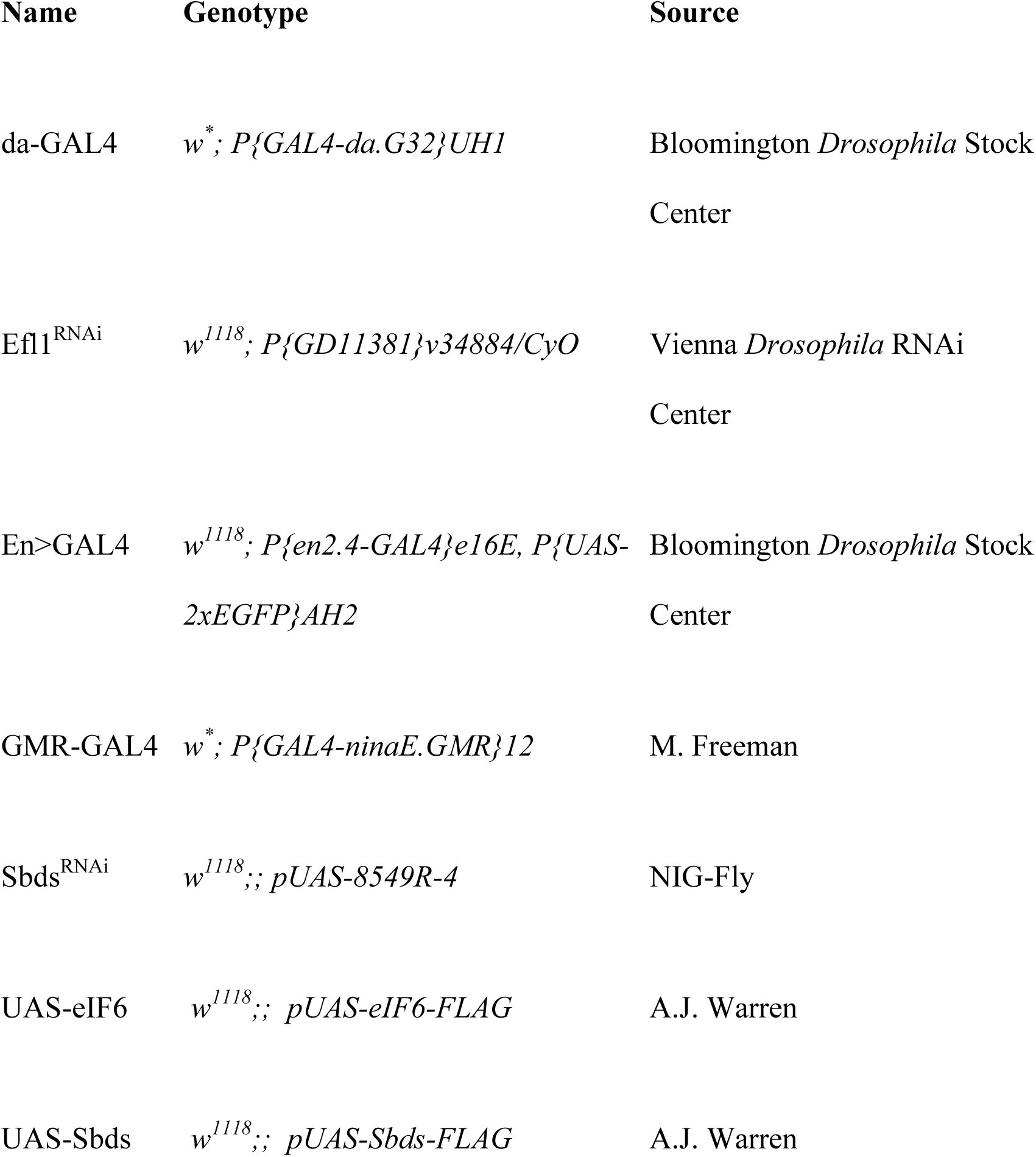
*Drosophila melanogaster* strains.

**Supplementary Table 5.**
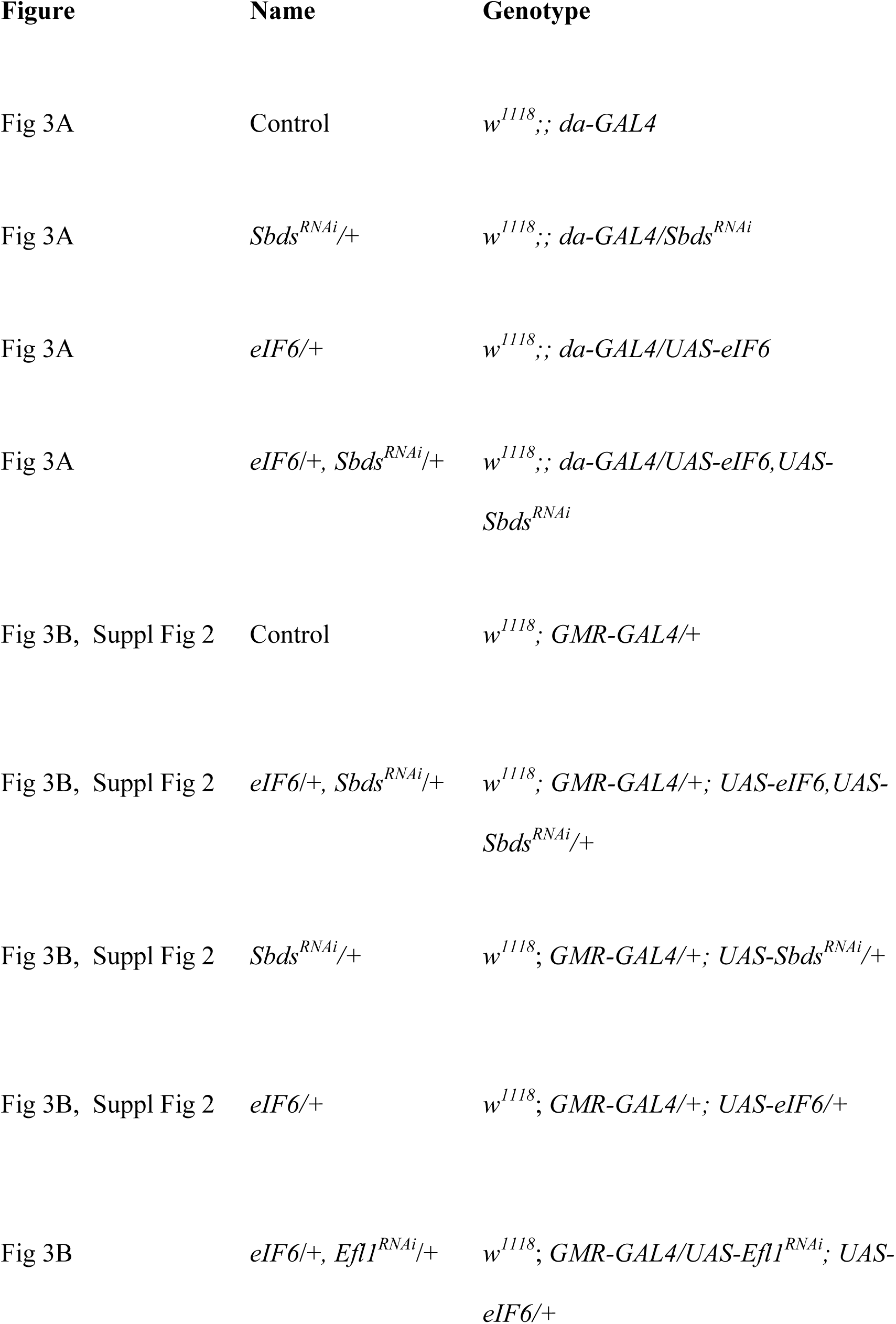
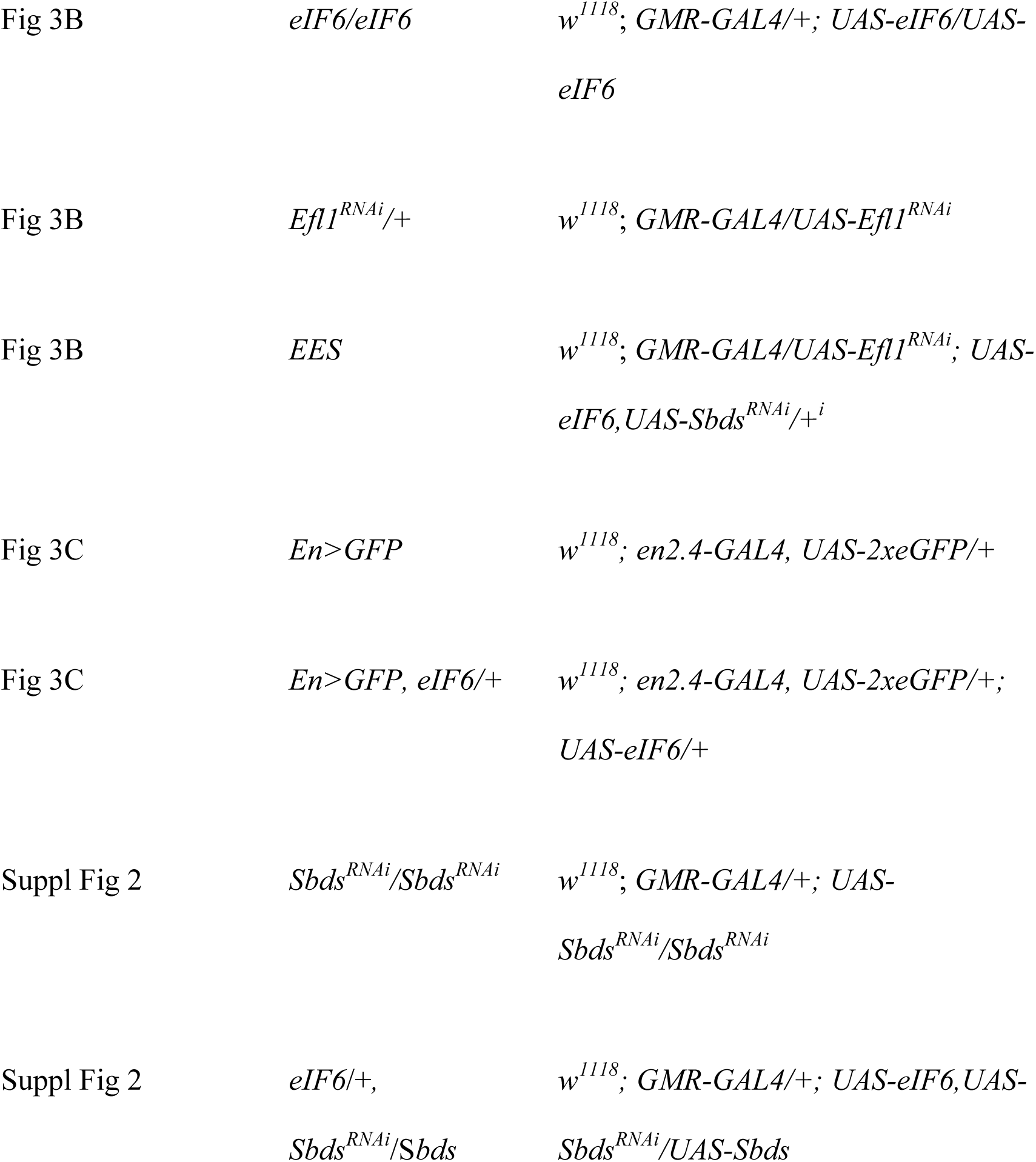
*Drosophila melanogaster* genotypes.

